# Single-cell multi-omic topic embedding reveals cell-type-specific and COVID-19 severity-related immune signatures

**DOI:** 10.1101/2023.01.31.526312

**Authors:** Manqi Zhou, Hao Zhang, Zilong Bai, Dylan Mann-Krzisnik, Fei Wang, Yue Li

## Abstract

The advent of single-cell multi-omics sequencing technology makes it possible for researchers to leverage multiple modalities for individual cells and explore cell heterogeneity. However, the high dimensional, discrete, and sparse nature of the data make the downstream analysis particularly challenging. Most of the existing computational methods for single-cell data analysis are either limited to single modality or lack flexibility and interpretability. In this study, we propose an interpretable deep learning method called multi-omic embedded topic model (moETM) to effectively perform integrative analysis of high-dimensional single-cell multimodal data. moETM integrates multiple omics data via a product-of-experts in the encoder for efficient variational inference and then employs multiple linear decoders to learn the multi-omic signatures of the gene regulatory programs. Through comprehensive experiments on public single-cell transcriptome and chromatin accessibility data (i.e., scRNA+scATAC), as well as scRNA and proteomic data (i.e., CITE-seq), moETM demonstrates superior performance compared with six state-of-the-art single-cell data analysis methods on seven publicly available datasets. By applying moETM to the scRNA+scATAC data in human bone marrow mononuclear cells (BMMCs), we identified sequence motifs corresponding to the transcription factors that regulate immune gene signatures. Applying moETM analysis to CITE-seq data from the COVID-19 patients revealed not only known immune cell-type-specific signatures but also composite multi-omic biomarkers of critical conditions due to COVID-19, thus providing insights from both biological and clinical perspectives.

## 1 Introduction

Multi-omic single-cell high-throughput sequencing technologies opens up new opportunities to interrogate cell-type-specific gene regulatory programs. Single-cell RNA sequencing (scRNA-seq) combined with Assay for Transposase-Accessible Chromatin using sequencing (ATAC-seq) [1] simultaneously measure the transcriptome and chromatin accessibility in the same cell. CITE-seq [2] measures surface protein and transcriptome data using oligonucleotide-labeled antibodies. By integrating the information from these multiple omics, we can expand our understanding of the genome regulation from multiple perspectives.

However, extracting meaningful biological patterns from the fast-growing multi-omic single-cell data remains a challenge due to several factors [3, 4]. Firstly, multi-omic single-cell technologies are still in the early stages. The cell yield is lower compared to the single-omic technologies such as scRNA-seq. On the other hand, the combined feature dimension is much higher (e.g., genes and peaks). This requires a more deliberate model design that can flexibly distill meaningful cell-type signatures from the multi-modal data while not overfitting the data. Secondly, multi-omic single-cell data are noisier compared with bulk-level or single-cell single-omic data. This calls for a probabilistic model that can infer latent cell types while properly accounting for the statistical uncertainty. Thirdly, the batch effects make it challenging to distinguish biological signals from study-specific confounders. Lastly, multi-omic single-cell are more costly compared to scRNA-seq or scATAC-seq alone. It is therefore highly cost-effective if we can profile single-omic data (e.g., transcriptome) and then predict the unobserved omic (e.g., chromatin accessibility or proteome). Nonetheless, the prediction from one modality to another is a challenging task, particularly from low dimension to high dimension.

Recently, several computational methods were developed to tackle the above multi-modality data integration challenges encountered in multi-omic single-cell data analysis. For instance, SMILE [5] integrates multi-omic data by minimizing the mutual information of the latent representations among the modalities and batches. The totalVI [6] and multiVI [7] integrate CITE-seq data and scRNA+scATAC data via variational autoencoder (VAE) frameworks, respectively. Cobolt [8] is a hierarchical Bayesian generative model to integrate cell modalities. scMM [9] is a mixture-of-experts (MoE) model developed to impute one missing modality conditioned on the other. Multigrate [10] adopted a product-of-experts (PoE) framework to integrate multi-omic data. MOFA+ [11] uses mean-field variational Bayes and coordinate ascent to fit a Bayesian Group Factor Analysis model to integrate the multi-omic data. Seurat V4 [12] integrated multimodal single-cell data through the weighted nearest neighbor algorithm. While many of these methods conferred promising performances in some of the tasks such as cell clustering or modality imputation, they often need to compromise scalability, interpretability, and/or flexibility.

In particular, when a neural network is used to encode the high-dimensional multi-omic data, interpretability is traded for flexibility; when a linear model or independent feature assumption is made, flexibility is traded for interpretability and scalability. However, all three are important to reveal cell-type-specific multi-omic signatures that are indicative of gene regulatory programs from large-scale data. Furthermore, most of these methods are entirely data-driven and therefore incapable of fully utilizing the existing biological information such as gene annotations or pathway information.

In this study, we present Multi-Omics Embedded Topic Model (moETM) to integrate multiple molecular modalities at the single-cell level. As one of the main technical contributions, moETM uses product-of-experts to infer latent topics underlying the single-cell multi-omic data and a set of linear decoders to learn shared embedding of topics and multi-omic features (e.g., genes, chromatin accessibility, and/or protein) that can accurately reconstruct the high-dimensional multi-omic data from their low-dimensional latent topic space (**Fig.**1a). Using stochastic amortized variational inference, moETM is highly scalable to large multi-omic datasets containing over 40,000 cells from scRNA+scATAC-seq and over 200,000 cells from CITE-seq data. Through effectively integrating multiple modalities from multi-omic single-cell sequencing data, moETM seeks to achieve 3 tasks: (1) clustering cells into biologically meaningful clusters to identify sub-celltype indicative of phenotype of interests (**Fig.** 1b); (2) imputing one omic (e.g., single-cell transcriptome or surface proteome) using the other omic (e.g., chromatin accessibility or single-cell transcriptome) (**Fig.** 1c); (3) identifying cell-type signatures, which serve as biomarkers for a target phenotype (**Fig.** 1d). Through comprehensive experiments on seven single-cell multi-omic datasets, we demonstrate moETM’s ability comparatively with six state-of-the-art (SOTA) computational methods. We further showcase how moETM facilitates the analysis of the COVID-19 single-cell CITE-seq dataset. Quantitatively, we observe that moETM learns the joint embeddings from multiple modalities with better or comparable bio-conservation, batch-effect correction, and cross-modality imputation compared with the existing methods [5–9, 11, 12]. Furthermore, the topic embedding learned by moETM can be used to gain biological insights into the cell-type-specific mutli-omic regulatory elements.

**Figure 1:**
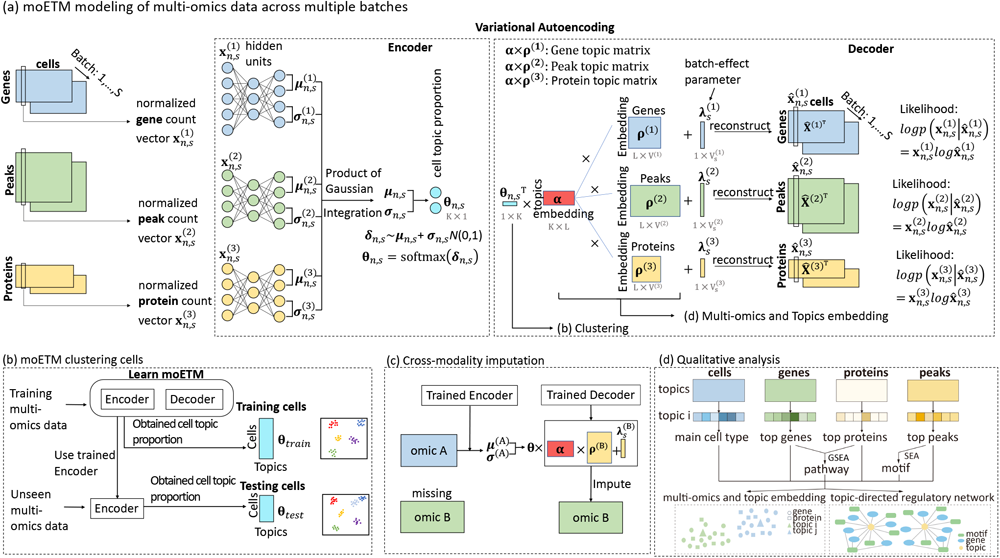
moETM model overview. **a**. Modeling single-cell multi-omics data across batches. In a nutshell, moETM integrates *M* omics via the product-of-experts (PoE), each of which is a pair of encoder and decoder. For a given cell *n* from batch *s*, each expert encoder takes one omic *m* as input **x**^(^*^m^*^)^ and produces the mean ***µ***^(^*^m^*^)^ and log variance log((***σ***^(^*^m^*^)^)^2^) for the omic-specific Gaussian distributed latent embedding variable. The product of these Gaussian densities over the *M* omics is also a Gaussian, from which we sample a joint logistic Gaussian latent embedding ***θ****_n,s_ ∼* softmax(***µ****_n,s_* + ***σ****_n,s_ N* (0, **I**)) to represent the cell. Each linear *m^th^* decoder expert then takes the same topic proportion ***θ****_n,s_* as input and reconstruct the original omic *m* for the cell with the aid of the global topic embedding ***α*** and the omic-specific feature embedding ***ρ***^(^*^m^*^)^. The end-to-end learning of the encoder network parameters and the decoder topic and feature embeddings is accomplished by maximizing the evidence lower bound of the categorical likelihood for the multi-omic count data via backpropagation. **b**. Evaluating moETM through cell clustering. The trained PoE encoders is used to infer the topic proportion of either training ***θ****_train_* or test data ***θ****_test_* from their multi-omic data. The integration performance of moETM is evaluated by clustering cells based on their topic proportion and qualitatively evaluated by UMAP visualization. **c**. Cross-omic imputation. To impute the missing omic *B* (e.g., protein) for a test cell, the trained moETM feeds the observed omic input vector **x**^(^*^A^*^)^ to the corresponding encoder expert *A*. The joint Gaussian embedding is then fed to the expert decoder *B*, which takes the inner product of the cell embeddings with its learned topic embedding and feature embedding for omic *B*. **d**. Downstream topic analysis. The learned topics-by-{cells, genes, proteins, peaks} matrices enable identifying cell-type-specific topics, gene signatures, surface protein signatures, and regulatory network motifs, respectively.

## 2 Results

### 2.1. moETM model overview

As an overview, moETM integrates multiomics data across different experiments or studies with interpretable latent embeddings (**Fig.** 1). It is built upon the widely used variational autoen-coder (VAE) [13] to model multi-modal data (**Fig.** 1a). However, to tailor the VAE framework for the single-cell multi-omic data, we made two main contributions on both the encoder and the decoder of the VAE.

The encoder in moETM is a two-layer fully-connected neural network, which infers topic proportion from multi-omic normalized count vectors for a cell. We assume the latent representation of each omic follows a *K*-dimensional independent logistic Normal distribution. Our goal is to effectively combine these distributions into a joint distribution of the multi-omic data. To this end, we take the product of the *K*-dimensional Gaussians i.e., Product-of-Gaussians (PoGs). Because the PoGs is also a Gaussian density function, we can represent the joint latent distribution in closed-form. In principle, this results in a tighter ELBO and therefore more efficient variational inference compared to the mixture of experts (MoE) approaches [14] as adopted in MultiVI/TotalVI [6, 7] and scMM [9]. In particular, these MoE approaches sample *K*-dimensional Gaussian variables for each omic and then take their average. In contrast, our PoG formalism requires sampling only once from the joint Gaussian. Therefore, moETM may confer more robust estimates thanks to the reduced sampling noise from the Monte Carlo approximation procedure. To obtain interpretable cell embedding, we perform a softmax transformation on the joint Gaussian density. The resulting logistic Normal distribution can be considered as a topic mixture membership for the cell. These topics can be directly mapped to known cell types based on their top gene signatures detected from our linear-decoder (as described below). Because the topic distribution must sum to 1 over the *K* topics, the inferred topic mixture membership of a cell express statistical uncertainty in the cell embedding.

On the decoder side, inherited from our earlier work [15], moETM employs a linear matrix factorization to reconstruct the normalized count vectors from the cell embedding. Our working hypothesis is that the encoder creates a linearly separable space for the decoder to achieve a good reconstruction when the two networks are trained end-to-end. Specifically, the decoder factorizes the cell-by-feature matrices into a shared cell-by-topic matrix **Θ**, a shared topic-embedding matrix ***α***, and *M* separate feature-embedding matrices ***ρ***^(^*^m^*^)^, where *m ∈ {*1*,. . ., M}* indexes the omics. Since different omics share the same cells-by-topics matrix but had their own feature-embedding matrices, we can explore the relations among cells, topics, and features in a highly interpretable way. This departs from the existing VAE models such as scMM [9], BABEL [16], and Multigrate [10] that used a neural network as the decoder. Another main challenge in single-cell data analysis is the batch effects, which are sources of technical variation. To account for those, we introduced the omic-specific batch-removal factors ***λ***^(^*^m^*^)^ *∈* R*^V^* ^(^*^m^*^)^*^×S^* for each omic *m* (**Fig.** 1a), which acts as a linear-additive batch-specific bias in reconstructing each modality. By regressing out the batch effects via ***λ***^(^*^·^*^)^, moETM can learn biologically meaningful representation in terms of the cell topic mixture and the topic/feature embedding. As detailed in **Methods**, all the parameters in moETM are learned end-to-end by maximizing a common objective function defined as the evidence lower bound (ELBO) of the marginal data likelihood under the framework of amortized variational inference.

### 2.2. Multiomics integration

We performed quantitative evaluations of moETM on the integrated low-dimensional representation comparing with six state-of-the-art multiomics integration methods (SMILE [5], scMM [9], Cobolt [8], MultiVI/TotalVI [6, 7], MOFA+ [11], Seurat V4 [12]) on seven published datasets. Four out of the seven datasets are single-cell transcriptome and chromatin accessibility (gene+peak) datasets and the other 3 are single-cell transcriptome and surface protein expression (gene/transcript+protein) datasets measured by Cellular Indexing of Transcriptomes and Epitopes by sequencing (CITE-seq).

The performance of the multiomics integrative task were based on both biological conservation metrics and batch removal metrics (**Methods**). For the biological conservation score, we adopted the common metrics including Adjusted Rand Index (ARI) [17] and Normalized Mutual Information (NMI [18]). For evaluating batch-effect removal, we used k-nearest-neighbor batch-effect test (kBET) [19] with graph connectivity (GC).

To make a comprehensive comparison, we used three experimental settings: *i*) 60/40 random split for training and testing with 500 repeats (**Table** 1 and 2, **Supplementary Table S1 and S2**); *ii*) training and testing both on the whole dataset (**Supplementary Table S3**); *iii*) training and testing across different batches (**Supplementary Table S4 and S5**). The number of topics was set to 100 during the training based on the robust performance (**Supplementary Fig. S1**). Overall, we obtained consistent results across all 3 settings and therefore chose to focus on describing the results based on the first setting.

We observed that moETM achieved the best overall performance when averaging over all datasets’ performance scores among 3 out of 4 evaluation metrics. Similarly, when averaging across gene+protein datasets only, moETM achieved the best overall performance among 3 out of 4 evaluation metrics (**Fig.** 2 middle panel). In particular, moETM conferred the highest averaged ARI, NMI, and GC when either averaging over all datasets or averaging over gene+protein datasets specifically, and it also conferred the second highest averaged kBET, which is only marginally behind multiVI/totalVI. The latter two methods might have over-corrected the batch effect at the expense of biological conservation. When averaging over gene+peak datasets, moETM can still achieve the best among 2 out of 4 evaluation metrics. Specifically, moETM ranked the second highest on ARI and NMI and slightly behind Seurat V4, which has a larger standard deviation compared with moETM.

**Figure 2:**
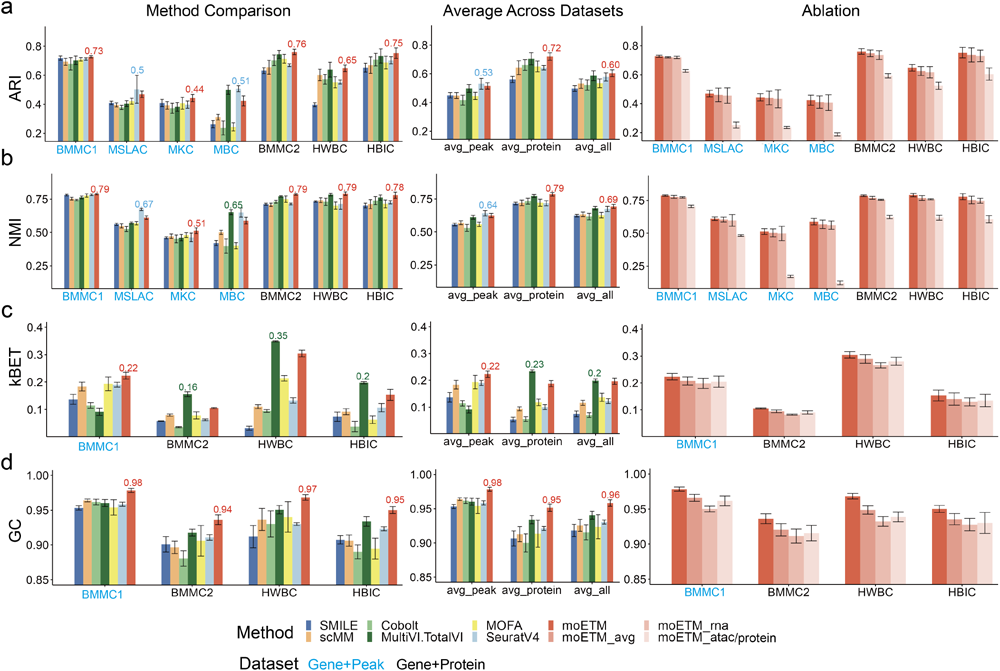
Methods comparison based on cell clustering. The left column illustrates the individual performance of each method on each dataset. The 7 datasets are indicated on the x-axis with gene+peak datasets colored in blue and gene+protein datasets colored in black. The evaluation scores for each are shown on y-axis. Ten colors were used to represent 10 different methods including six existing state-of-the-art methods, the proposed moETM model, and 3 of its ablated versions. Within each dataset, the highest value was labeled on the top of the corresponding bar. The middle column is the comparison of averaging values across datasets for each method. The right column is the comparison between moETM and its three ablated versions. Each row represents an evaluation metric. **a**. Adjusted Rand Index (ARI). **b**. Normalized Mutual Information (NMI). **c**. k-nearest neighbour batch effect test (kBET). **d**. Graph connectivity (GC).

For individual datasets, moETM is either the best or the second best method on 6 out of 7 datasets (except MBC) for different experimental settings in terms of the ARI (**Fig.** 2a,**Table** 1 and 2). One possible reason could be that the sample size of MBC (3293 cells) from which moETM learns high-dimensional peak embeddings is small compared with the other 6 datasets. To assess the benefits of the added features in moETM, we compared moETM with its ablated versions: moETM_rna, moETM_atac, and moETM_protein, where moETM was trained on a single omic. As expected, the performance of moETM on single modality decreased, indicating that moETM could improve its performance by leveraging multiple modalities (**Fig.** 2 right panel).

Similar quantitative conclusions can be drawn based on NMI (**Fig.** 2b, **Supplementary Table S1**, and **Supplementary Table S2**). For kBET (**Fig.** 2c), moETM is the best for the BMMC1 dataset and the second best on BMMC2, HWBC, and HBIC datasets – slightly behind MultiVI/TotalVI. Therefore, while moETM conferred higher biological conservation scores in terms of ARI and NMI, it still maintains a comparable kBET scores on all four datasets compared to MultiVI/TotalVI. Indeed, we observed an excellent balance between the biological conservation and batch effect removal because moETM achieved notably higher GC compared to all methods (**Fig.** 2d). This is because GC is the only metric that is based on both the cell types and batch labels by measuring the similarity among cells of the same type from different batches based on the embedding learned by each method [20].

We postulated that the main reason for the moETM’s superior integration performance is it’s the Product of Gaussians (PoG) formulation. To that end, we constructed moETM_avg, which replaced PoG with averaging of sampled variables from individual Gaussian distributions similar to the existing VAE models like scMM [9]. As expected, the performance of moETM_avg was worse than moETM in all datasets in terms of both bio-conservation and batch removal evaluation metrics (**Fig.** 2 right panel). Furthermore, since scMM also adopted the average of the Gaussian samples from the encoder, the fact that moETM_avg outperformed scMM indicates the benefits of using the linear decoder, which further improves the multi-omic integration while correcting batch effects across all cells.

We further verified the clustering performance by visualizing the cell topic mixture embeddings using Uniform Manifold Approximation and Projection (UMAP) [21] (**Fig.** 3). Indeed, not only did moETM remove batch effects but also revealed a better representation of cell type clusters. For example, “Plasmablast IGKC-” cells were grouped closely by moETM but were clustered into multiple small parts by SMILE (**Fig.** 3a). Moreover, plasmablast cells from different batches were also mixed better by moETM compared with SMILE, which indicated a better batch-effects correction. “CD4+ T activated” and “CD4+ naive” cells were closer within the same cluster but clearly distinguishable between themselves. In contrast, these two cell types were mixed together by SMILE and scMM. In modeling the BMMC1 dataset (gene+peak), “B1 B” cells and “naive CD20+ B” cells (**Fig.** 3b) were mixed by other methods while better separated by moETM. In addition, we visually compared the cell clustering by the individual modalities with that by the integrated modalities via UMAP (**Fig.** S6). Specifically, utilizing the BMMC1 dataset, we used the encoders for the individual omics to generate the cell topic mixtures separately for the RNA and chromatin accessibility (i.e., peaks) and compared them with the cell topic mixtures after integrating the two omics via PoG. Similarly, we also compared the cell clustering based on the individual omics of single-cell transcriptome and surface proteins with the clustering based on the integrated RNA+protein topic mixture using the BMMC2 dataset. We observed that the integrated topic representation led to a more coherent cell clustering compared with the topic mixture of individual omics. For example, the ‘CD14+ Mono’ cells were grouped more closely by the integrated topic mixture compared with the counterparts (**Fig.** S6a; ARI: 0.735 by RNA+peak in contrast to 0.690 by RNA only and 0.648 by peak only). Similarly, the “Plasmablast IGKC-” cells also formed tighter cluster in the integrated RNA+protein embedding space (**Fig.** S6b; ARI: 0.734 by RNA+protein in contrast to 0.688 by transcriptome only and 0.590 by surface proteins only). Therefore, moETM was able to improve cell clustering by integrating multiple modalities. Taken together, these results show that moETM is able to distinguish similar cell types by capturing biological information in its encoding space while removing batch effects.

**Figure 3:**
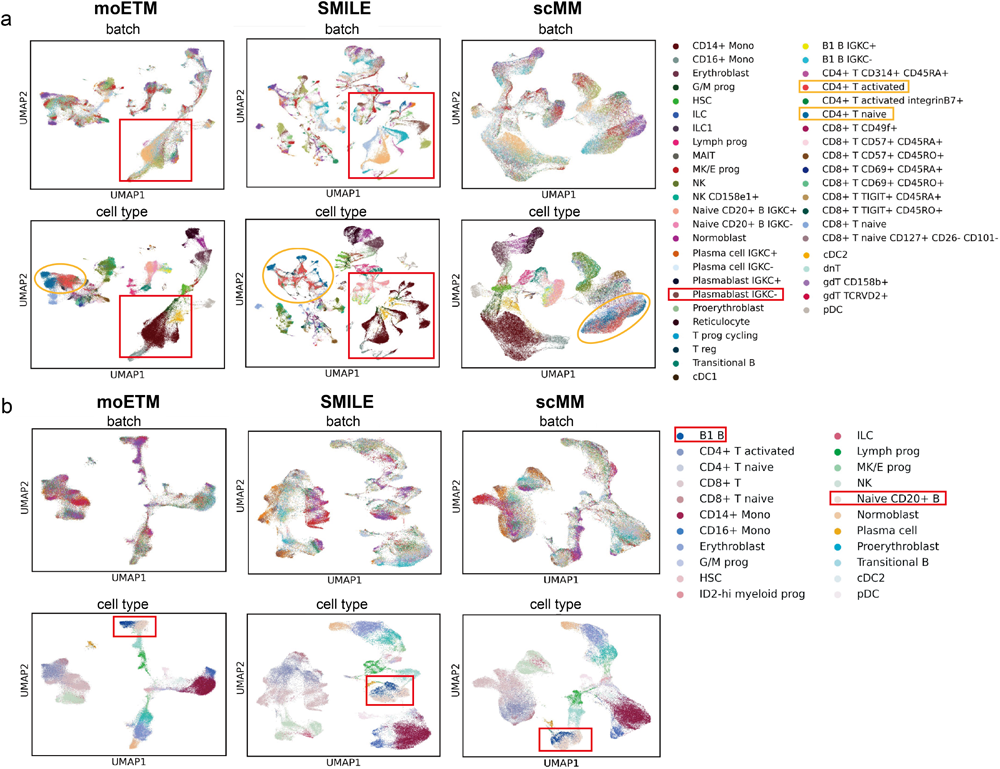
UMAP visualization of cell clustering. **a**. UMAP visualization of moETM, SMILE, and scMM on single-cell CITE-seq from BMMC2 dataset. Each point on the two-dimensional UMAP plots represents a cell. In the upper panel, different colors indicate different batches. In the lower panel, different colors indicate different cell types. **b**. UMAP visualization of moETM, SMILE, and scMM on the gene+peak multiome data from the BMMC1 dataset. Similarly to panel a, the upper and lower panel labelled with batch indices and cell types, respectively. The highlighted clusters and cell types in the legend were described in the main text.

### 2.3. Cross-omic imputation

In the case of gene+protein, moETM accurately imputes surface protein expression from gene expression for the BMMC2 dataset, achieving average Pearson (Spearman) correlation of 0.95, 0.92, and 0.88 (0.94, 0.90, and 0.85) on random split, leave-one-batch, and leave-one-cell-type imputation experiments, respectively (**Supplementary Table S6**). We visualized the reconstructed protein expression against the observed values using the BMMC2 (gene+protein) dataset (**Fig.** 4a). The imputed protein expression is highly linearly correlated with the observed one (**Fig.** 4b), which is what we expected given the high Pearson correlation of 0.95. The runner up methods - namely, scMM and BABEL - also performed well on this task, both achieving a correlation score of 0.94.

**Figure 4:**
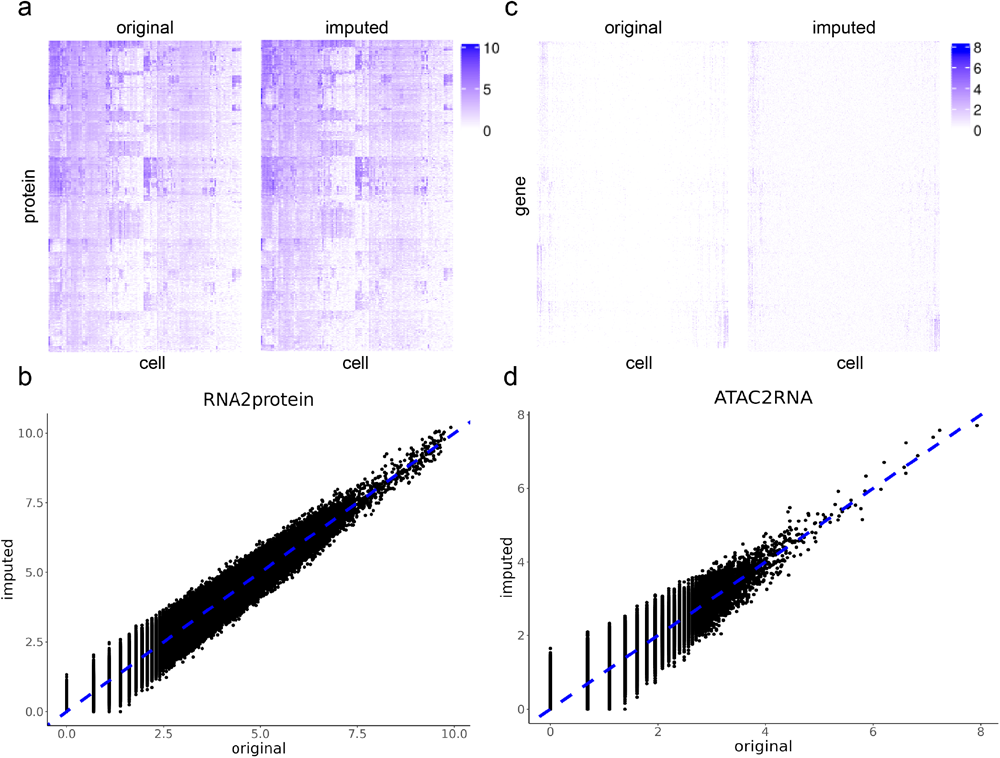
Cross-omic imputation. **a**. Heatmap of original protein and imputed protein values from gene expression using the BMMC2 CITE-seq dataset. We trained moETM on 60% of the cells with observed protein+gene omics and used the trained moETM to impute the protein expression based on the gene expression for the remaining 40% of the test cells. The two heatmaps correspond to the original and imputed protein expression, respectively. The columns are the randomly sampled 5000 test cells, and the rows are the surface proteins. For visual comparison, the column and row orders are the same for the two heatmaps. The color intensities are proportional to original or imputed protein expression over the cells. **b**. Scatter plot of original and imputed surface protein expression. The same values shown in panel a were displayed as scatter plot in this panel. The x-axis and y-axis represent the original and imputed protein expression values of the test cells, respectively. The diagonal line is in blue color. The more similar the reconstructed value is with the original value, the closer it is with the blue line, **c** & **d**. Heatmap and scatterplot of the original and imputed gene expression from chromatin accessibility on the BMMC1 dataset. The imputation results were shown in the same way as in panel a and c. We trained moETM on 60% of the cells with observed gene+peak omics. We then applied the trained moETM to the 40% test cells by imputing their gene expression based on their open chromatin regions (i.e., peaks). The original and imputed gene expression of the test cells were compared qualitatively in the heatmap and scatterplot. We also illustrated the imputation results from the low dimensional omic to the high dimensional omic in **Supplementary Fig. S2**.

Compared to the surface protein imputation task, imputing gene expression from the open chromatin regions is a more challenging task because of the sparser input scATAC-seq signals and the dynamic and often asynchronous interplay between the chromatin states and the transcriptome [22–24]. Nonetheless, moETM achieved a relatively high Pearson (and Spearman) correlation scores of 0.69, 0.65, and 0.58 (and 0.37, 0.35, and 0.32) on random split, leave-one-batch, and leave-one-cell-type experiments. These are notably higher than the corresponding correlation obtained by BABEL (Pearson: 0.65, 0.60, 0.55; Spearman: 0.34, 0.33, 0.30) and scMM (Pearson: 0.63, 0.61, 0.54; Spearman: 0.33, 0.33, 0.28) (**Supplementary Table S7**). Qualitatively, the imputed and the observed gene expression profiles also exhibit similar pattern and linear relationship (**Fig.** 4c, d).

In the previous two imputation applications, low dimensional modalities were generated from high dimensional modalities. The imputation from the low dimension to the high dimension is more difficult but nonetheless feasible. Specifically, on the 3 same experimental designs, the Pearson (and Spearman) correlations between the observed and the imputed open chromatin regions from gene expression are 0.58, 0.55, and 0.51 (and 0.33, 0.30, and 0.28) (**Supplementary Table S8**); the Pearson (and Spearman) correlation between the observed and imputed gene expression from protein expression are 0.65, 0.63, and 0.60 (and 0.41, 0.39, and 0.37) (**Supplementary Table S9**). In contrast, the runner-up method scMM achieved Pearson (and Spearman) correlations of 0.40, 0.29, and 0.37 (and 0.29, 0.25, and 0.21). For imputing chromatin accessibility from gene expression. For imputing gene expression from surface protein, scMM and BABEL also fell behind moETM in terms of both Pearson and Spearman correlations (**Supplementary Table S9**). Qualitatively, the imputed and the observed peaks and gene expression exhibit consistent patterns (**Supplementary Fig. S2a, c**) and strong linear trends and similar patterns (**Supplementary Fig. S2b, d**).

### 2.4. Correlating RNA transcripts with surface proteins and in-cis chromatin accessibility regions

As a proof-of-concept, we sought to assess whether the top surface proteins can be mapped to the top genes under the same topic (i.e., following the central dogma). To this end, we trained a 100-topic moETM on the BMMC2 (gene+protein) dataset generated by CITE-seq over 90,000 cells. For each topic, we calculated the Spearman correlation of topic scores between the 134 pairs of the gene transcripts and the corresponding translated surface proteins (**Fig.** 5a). The correlations ranged from -0.096 to 0.751 with an average of 0.29. In particular, 96 of the 100 topics have positive correlations. Among them, 13 topics have correlations larger than 0.5. In particular, the correlation in topic 40 was 0.576, and the correlation in topic 44 was 0.628.

**Figure 5:**
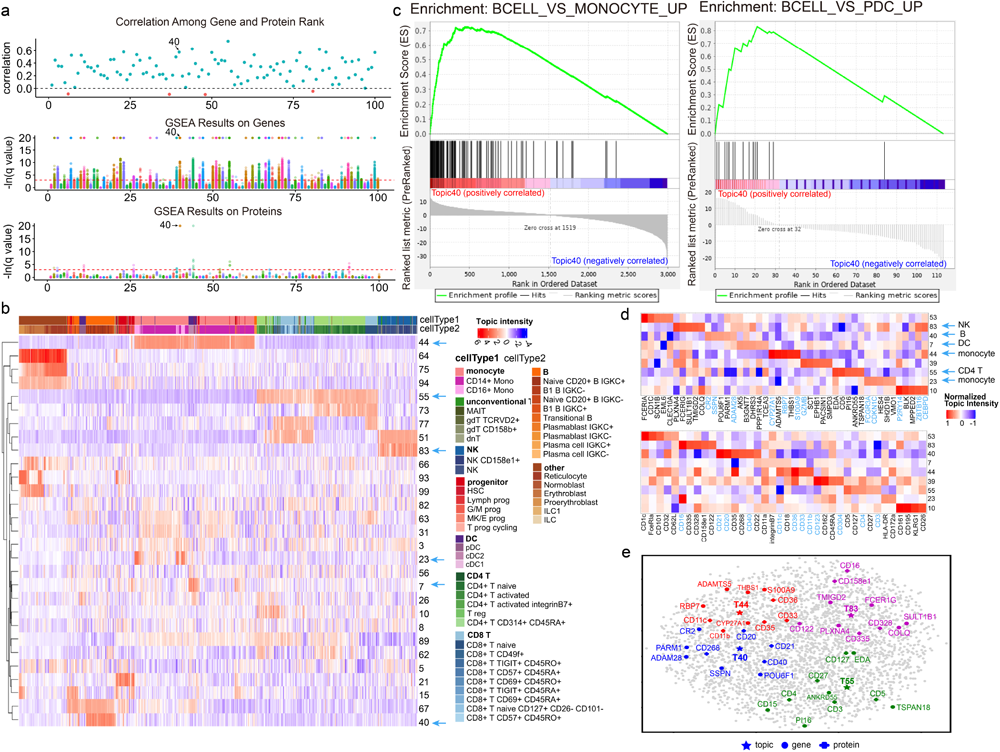
Topic analysis of gene+protein CITE-seq data. **a**. Protein-RNA correlations and pathway enrichments for the 100 topics learned from the CITE-seq BMMC2 data. In each plot, the x-axis is the 100 topics and the y-axis is either the protein-RNA correlation or the pathway enrichment scores in terms of -ln q-value. The top panel is the Spearman correlation between the RNA and protein expression for the same genes under each topic. Correlations above 0 are labeled blue and correlations below 0 are labeled red. The middle and the bottom panels are the corresponding GSEA enrichments of gene and protein topic scores, respectively. The dots correspond to the tested immunologic signature gene sets from MSigDB. Different colors represent different gene sets. **b**. Topics embedding of 10,000 sub-sampled cells from the BMMC2 dataset. Only the topics (rows) with the sum of absolute values greater than the third quartile across all sampled cells (columns) were shown. The two color bars display two tiers of annotations for the 9 broad cell types (cellType1) and 45 fine-grained cell types (cellType2). The topics that were labelled with arrows were described in the main text. **c**. GSEA leading-edge analysis of Topic 40. The left panel is the GSEA result of gene topic scores on a significantly enriched gene set (q-value *<* 0.001), which contains up-regulated genes in B cells relative to the monocytes. Similarly, the right panel displays an enriched gene set (q-value *<* 0.001), based on the protein topic scores for the same topic. The gene set contains up-regulated genes in B cells relative to plasmacytoid dendritic cells (pDC). **d**. Genes and proteins signatures of the select topics. The upper and lower panels display the topics-by-genes and topics-by-proteins heatmap, respectively. The top genes and proteins that are known cell-type markers based on CellMarker or literature search are highlighted in blue. For visualization purposes, we divided the topic values by the maximum absolute value within the same topic such that the topic scores range between -1 and 1. **e**. UMAP visualization of the genes, proteins, and topics via their shared embedding space. Genes, proteins, and topics were labeled by star, circle, and cross shapes respectively. Topics 40, 44, 55, and 83 were colored in blue, red, green, and purple respectively. The corresponding topic indices and gene/protein symbols were highlighted by corresponding colors.

To further quantify the known transcript-protein as well as gene-peak regulatory relations, we computed their Spearman correlations across topics inferred from the BMMC datasets. We paired a peak with a gene if it is within 150K bp distance from the transcription start site (TSS) of the gene. The overall distribution of the correlations for transcript-protein pairs and gene-peak pairs were both significantly higher than 0 (p *<* 2.2e-16; one sample t-test) and comparable to the correlations calculated directly from the observed data (**Supplementary Fig. S8a,c**). Specifically, the transcript-protein correlations ranged from -0.265 to 0.642, with an average of 0.245; the gene-peak correlations ranged from -0.09 to 0.67, with an average of 0.21.

Notably, 90% transcript-protein pairs exhibited positive correlations and 9 pairs displayed correlations exceeding 0.5 (**Supplementary Fig. S8e**). Nonetheless, several transcript-protein pairs exhibited low or negative correlations. Several factors could contribute to these low correlations. Firstly, random noise may hinder correlations between genes and proteins. Secondly, dynamic cellular processes at the single-cell level can give rise to variations between cells, leading to a decrease in correlations [25]. For example, transcriptional bursting or delays between transcription and translation will lead to asynchronous behavior of gene and protein during the cell cycle, thereby reducing the correlations between gene and protein expression levels [25]. Particularly, a number of transcript-protein pairs displayed negative correlations (**Supplementary Fig. S8e**). This phenomenon has also been observed in previous studies [26–28]. For instance, Li et al. (2020) reported a mismatch between mRNA and protein expression levels, including the *CD69*-CD69 pair [27]. One possible cause might be due to the impact of other biological processes overriding the effects of transcription [28]. Taking *CD69*-CD69 as an example, the *CD69* gene may undergo post-translational modifications such as differential glycosylation [29]. The transcribed *CD69* mRNA molecules can be translated to 22.5 kDa polypeptide, which can further be differentially glycosylated to two different subunits. These subunits can be randomly combined to form different receptors, leading to a reduction in the abundance of the CD69 protein [29]. If the influence of post-translational modifications surpasses the impact of protein synthesis, it can give rise to a negative correlation. Furthermore, the *CD69* mRNA transcripts are unstable. The level of *CD69* mRNA could decline rapidly with a half-life of less than 60 minutes [29]. While mRNA molecules degrade over time, protein levels may maintain relatively stable or continue to accumulate. If the rate of mRNA degradation surpasses that of protein synthesis, a negative correlation could emerge.

In addition to investigating correlations across topics, we calculated correlations across cells by computing Spearman correlations in terms of observed values and reconstructed values based on the BMMC datasets (**Supplementary Fig. S8a-d**). The correlations based on both the observed and reconstructed data across cells were significantly greater than 0, indicating consistent relations among transcript-protein and gene-peak pairs captured at the cell level. However, the correlations from the reconstructed values are higher than those from the observed values (**Supplementary Fig. S8b,d**). This is because the observed values may contain random noise or batch effects compared with reconstructed values by moETM, which can be considered as denoised and confounder-corrected values of the gene/protein/peak signals.

### 2.5. Immune cell-type signatures revealed by multi-omic topics learned from CITE-seq data

To identify cell-type signatures, we associated each topic with the specific cell type that exhibit the highest average topic score across cells. Notably, not all topics were uniquely associated with one single cell type and some topics might be enriched for a combination of multiple cell types. Therefore, we chose to describe a selected subset of the topics based on their distinctly enriched cell types and heatmap visualization (**Fig.** 5b). For instances, topic 44 was associated with monocytes, which consists of CD14+ and CD16+ Mono; topic 40 was associated with B cells, which consists of primarily Naive CD20+ B IGKC+ and Naive CD20+ B IGKC-cells; topic 83 was associated with natural killer cells. These are visually easy to detect from the topic mixture probabilities among the individual cells (**Fig.** 5b).

Under each cell-type-enriched topic, many top genes and top proteins are the known cell-type markers (**Fig.** 5d). For example, under topic 40 (i.e., a B-cell topic), the top genes *CR2*, *SSPN*, and *ADAM28* are known marker genes for B cell; the top proteins CD21, CD20, and CD40 are also marker proteins for B cells according to the CellMarker database [30]. For topic 7, one of top proteins CD11c is a marker protein for dendritic-cell [30]. For topic 83, protein CD16, marker for natural killer cells, is among its top proteins [30]. For topic 44, the top gene *S100A9*’s coding protein is a chemotactic factor for monocytes [31] and is highly expressed during inflammatory processes [32]; among the top proteins for topic 44, CD36 [33], CD33, and CD11c [34] are also markers for monocyte sub-cell-types. Similarly, monocyte is also enriched in topic 23, which shares the top marker protein CD16 with topic 44 but also contains unique top genes such as *CDKN1C* and *FCGR3A*. While *CDKN1C* is a known marker gene for monocyte [35], *FCGR3A* is up-regulated in CD16+ monocytes as supported by the existing literature [36].

Moreover, we performed Gene Set Enrichment Analysis (GSEA) [37, 38] using the topic scores for all of the genes and proteins. Because BMMCs are immune cells, we queried the C7 ImmuneSigDB from MSigDb, which is a collection of 5219 gene sets related to immune pathways [39–41]. Across all 100 topics, we identified 2569 enriched gene sets with q-value *<* 0.05 using gene topic scores and 22 enriched gene sets using protein topic scores (**Fig.** 5a). For example, in topic 40, using the gene topic scores, we found a gene set that consists of up-regulated genes in B cells with respect to monocytes [42] (**Fig.** 5c left panel); using the protein topic scores, we found a gene set that consists of up-regulated genes in B cells compared to plasmacytoid dendritic cells (PDC) (**Fig.** 5c right panel) [42].

Furthermore, we projected the topic embeddings and feature embeddings onto a common 2D space using UMAP (**Fig.** 5e). We observed that the top marker genes and the top marker proteins for the cell type clustered together around the corresponding topics, implying a well aligned embedding space within and across these modalities. Together, the results suggested that the cell-type-enriched topics inferred by moETM from the CITE-seq data reveal meaningful biological relations between genes and proteins.

### 2.6. Joint multi-omic topic analysis identified cell-type-specific pathways and regulatory motifs

The topic embedding learned from the scRNA+scATAC data enables us to investigate the relationship between top genes and top peaks (i.e., top open chromatin regions) in the cell-type-specific topics. Given that many top genes are known cell-type markers (**Fig.** 6a), we postulated that the top peaks could be associated with the top genes via *in-cis* or *in-trans* regulatory elements. Different from the previous gene+protein case, one challenge in interpreting the gene+peak multiomic topics is that peaks cannot be matched directly with genes. We proposed two approaches to solve this issue. One is to link peaks to their nearby genes to obtain the peak-neighboring-genes (**Methods**). The other approach is to identify enriched motifs among the top peaks and explore the relationship between genes and motifs via the corresponding transcription factors (TFs) and their target genes.

**Figure 6:**
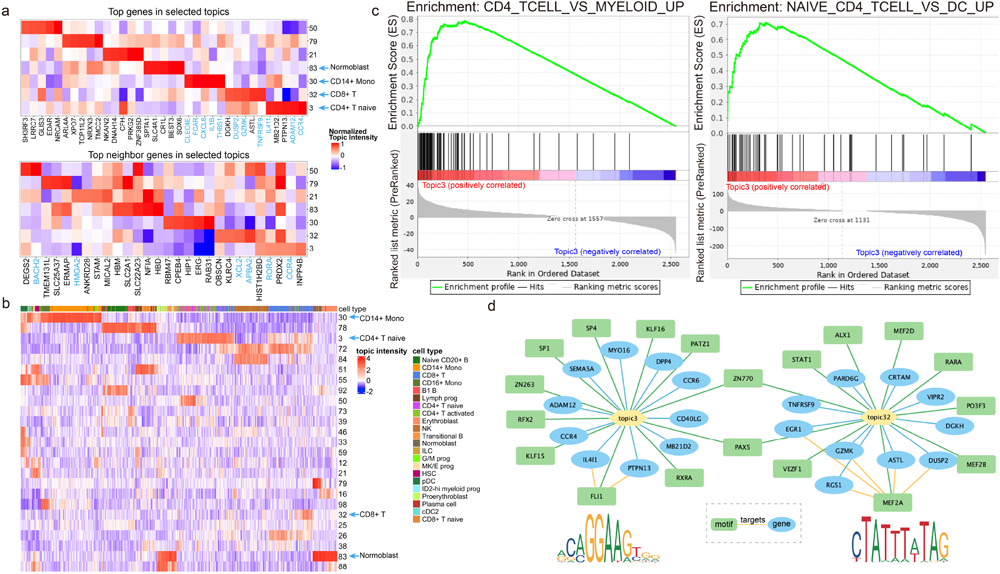
Topic analysis of single-cell gene+peak data from the BMMC1 dataset. **a**. Top genes and top peak-neighbour-genes of the select topics. The heatmap displays the top features (columns) for 7 out of 100 topics, which were selected based on their cell-type enrichments. The top signatures that are related to the enriched cell types based on CellMarker or literature search are highlighted in blue. For visualization purposes, we divided the topic values by the maximum absolute value within the same topic such that the topic scores range between -1 and 1. **b**. Topic embedding of cells from the BMMC1 dataset. The heatmap displays the embedding profiles of topics (rows) for 10,000 randomly sampled cells (columns) from the BMMC1 dataset. Only the topics with the sum of absolute values larger than the third quartile over the 10K cells are shown. The color bar on the top of the heatmap indicate the cell types with the text annotations shown in the legend. The columns and rows were ordered based on agglomerative hierarchical clustering with Euclidean distance and complete linkage **c**. GSEA leading edge analysis of Topic 3. The left panel is the GSEA result using gene topic scores and the right panel is the GSEA result using peak-neighboring-gene topic scores. The barcode in the middle are the genes that belong to the corresponding gene sets, namely the up-regulated genes in CD4 T cell relative to the Myeloid cells and the up-regulated genes in CD4 T cells relative to dendritic cells for the gene and peak modalities of the same topic, respectively. **d**. Topic-directed regulatory networks based on motif enrichment analysis. The blue ellipses represent genes and the green rectangles represent enriched motifs. The bottom left and right motif logos correspond to the transcription factors (TFs) FLI1 and MEF2A, respectively. The yellow edges between motifs and genes indicate known TF-target associations based on ENCODE TF Targets dataset.

For the first approach, the top genes and top peak-neighboring-genes in the select topics served as markers for the cell-type-specific gene regulatory programs (**Fig.** 6a). For example, topic 32 is associated with CD8+ T cell (**Fig.** 6a, b). We zoomed in the topic by examining its top genes and top peaks. <u>Three</u> of the top 5 genes (*<u>TNFRSF9</u>*, *ASTL*, *<u>GZMK</u>*, *<u>DUSP2</u>*, *DGKH*) were related to T cell. In particular, *GZMK* is a marker gene for T cells based on CellMarker [30]; *TNFRSF9* codes for a signaling protein that promotes expression of cytokines in CD8+ T cells [43]; *DUSP2* encodes an inducible nuclear protein and is highly expressed in T cells [44]. Among the top 5 peak-neighboring-genes (*<u>APBA2</u>*, *PRDX2*, *KLRC4*, *OBSCN*, *<u>XCL2</u>*), *APBA2* is a marker genes for cytotoxic CD8+ T lymphocyte [45]; *XCL2* expression levels substantially increased in CD8+ T cells during T cell activation [46].

As another example, topic 3 is associated with CD4+ T naive cells. <u>Three</u> out of the top 5 genes (*<u>CCR4</u>*, *<u>ADAM12</u>*, *PTPN13*, *MB21D2*, *<u>IL4I1</u>*) and <u>two</u> out of the top 5 peak-neighboring-genes (*INPP4B*, *<u>CCR4</u>*, *PRDX2*, *<u>RORA</u>*, *HIST1H2BD*) are related to T cells. Indeed, *CCR4* is shown to be specifically expressed among naive CD4+ T cells [47]; *ADAM12* is expressed in T cells in the inflamed brain and is a potential target for the treatment of Th1-mediated diseases [48]; *IL4I1* increases the threshold of T-cell activation and partially modulates CD4 T-cell differentiation [49]. For top peak-neighboring-genes, *RORA* is up-regulated among the activated CD4+ T cells [50].

To gain further mechanistic understanding of the inferred topics, we performed GSEA on the topic scores for the genes from the transcriptome modality and the topic scores for the peak-neighboring-genes from the chromatin-accessibility modality (**Supplementary Fig. S9**). Many enriched gene sets are related to the topic-associated cell types. For topic 3, for instance, one of the enriched gene sets based on the gene topic scores is up-regulated in healthy CD4 T cells with respect to healthy myeloid cells [51] (**Fig.** 6c). This is consistent to an enriched gene set from the peak-neighboring-gene analysis of topic 3, where the gene set consists of a set of genes that were up-regulated in naive CD4 T cell relative to the dendritic cells (DC) [52]. Therefore, GSEA further confirmed the cell-type-specific functions of the top genes and peak-neighboring-genes identified via moETM’s topics. Interestingly, the top transcripts and the top peak-neighbouring-genes do not often correspond to the same genes. This implies that the peaks and genes provide complementary information to (sometimes the same) cell-type-specific regulatory programs. Therefore, by effectively integrating the scRNA-seq and scATAC-seq data, the inferred multi-omic topics can reveal functional convergence at the pathway level.

Besides using peak-neighboring-genes, as the second approach, we also performed motif enrichment analysis on the top 100 peaks per topic (**Methods** 4.7, **Supplementary Fig. S9c**). We then constructed a putative regulatory network by linking the top genes and the enriched motifs via their associated topics (**Fig.** 6d). Interestingly, some of the top genes harbor those enriched motifs, implying that these genes are the putative target genes of the cognate TF. In topic 3, for example, one of the enriched motifs corresponds to a TF named FLI1 (p-value = 0.00117), and the top genes *IL4l1* and *PTPN13* are target genes of FLI1 based on the ENCODE Transcription Factor Targets [53, 54]. As another example, one of the enriched motifs for topic 32 corresponds to TF MEF2A (p-value = 5.21e-5), whose target genes include the top genes *RGS1*, *EGR1*, *GZMK*, *ASTL*, and *DUSP2* [53, 54].

We further expanded our topic-network analysis by including enriched pathways and cell types information (**Supplementary Fig. S7**). Specifically, we treated the topics, top genes, and enriched motifs/pathways/cell types as nodes, and interactions between topic-cell type, gene-motif, and gene-pathway were treated as edges. We defined the *intra-connections* within the same topic as edges between the topic nodes and cell type nodes. We also established *inter-connections* between genes and external nodes including motifs and pathways. Specifically, the top genes under each topic could serve as members of enriched pathways or target genes for enriched motifs. For instance, in topic 32, gene *DUSP2* is a target gene for enriched motifs MEF2A (p-value = 5.21e-5, Permutation test) and PAX5 (p-value = 1.53e-5, Permutation test), while also being a member gene in four enriched pathways and <u>three</u> of them are upregulated gene sets in the enriched cell type of T cells (<u>UNSTIM_VS_ACD3_ACD28_STIM_WT_CD4_TCELL_DN</u>, UNSTIM_VS_LPS_AND_ANTI_CD40_STIM_NIK_NFKB2_KO_DC_DN, <u>NKT_CELL_VS_ALPHAALPHA_CD8_TCELL_DN</u>, <u>UNSTIM_VS_ACD3_ACD28_STIM_WT_CD4_TCELL_DN</u>). Similarly, in topic 30, gene *FCAR* is a target gene for one enriched motif CEBPB (p-value = 1.27e-5, Permutation test) and a member of three enriched pathways and <u>two</u> of them are related to gene sets up-regulated in the enriched cell type of monocyte (<u>MONOCYTE_VS_MDC_UP</u>, PBMC_MRKAD5_HIV_1_GAG_POL_NEF_AGE_20_50YO_1DY_UP, <u>MONOCYTE_VS_MDC_DAY7_FLU_VACCINE_UP</u>). Likewise, in topic 3, gene *DPP4* is a target gene for two enriched motifs PAX5 (p-value = 5.39e-4, Permutation test) and SP1 (p-value = 1.13e-4, Permutation test) and a member of 11 enriched pathways where <u>four</u> of them (<u>NAIVE_TCELL_VS_NKCELL_UP</u> <u>NAIVE_TCELL_VS_MONOCYTE_UP</u>, <u>NAIVE_CD4_TCELL_VS_MONOCYTE_UP</u>, <u>NAIVE_CD4_TCELL_VS_DC_UP</u>) are up-regulated gene sets in CD4+ T naive cell. Those connections highlighted a consistent regulatory relationship across motifs and pathways under inferred topics.

Therefore, our multi-omic topic analysis suggests that some of the cell-type-specific regulatory programs are implicated with the sequence motifs and pathways. Further investigation is needed to establish the hierarchical relation between the TF and the cell lineage.

### 2.7. Prior pathway-informed enrichment

The single-cell multi-omic data are high-dimensional, sparse, and noisy. This is especially the case for the scRNA+scATAC-seq data because of the large number of genes and open chromatin regions. One way to further improve the interpretability of the topics derived from these data is by incorporating prior knowledge such as gene sets or pathway information. In the context of our moETM, this was done by fixing the embeddings-by-genes parameters to the observed pathways-by-genes matrix (**Methods**). Using the *∼*7000 Gene Ontology Biological Process (GO-BP) terms as the pathways-by-genes matrix, we trained the pathway-informed moETM (p-moETM) on the BMMC1 gene+peak dataset.

Quantitatively, p-moETM can achieve comparable cell-clustering performance with ARI of 0.72, which is only slightly lower than the default moETM that learned the gene embedding directly from the data (Table 1). We also identified several cell-types-specific topics along with their top genes and peaks (**Supplementary Fig. S3a-c**). Notably, the learned topics-by-embeddings matrix ***α*** from p-moETM are essentially the topics-by-pathways matrix. This allows us to directly identify the top pathways for each topic without performing post-hoc GSEA. For instance, topic 25 is associated with B1 B cell (**Supplementary Fig. S3a**). One of its top pathways is related to B cell activation (**Supplementary Fig. S3d**). As another example, topic 8 was enriched for the CD4+ T activated cell, and one of its top pathways was connected to the T cell apoptotic process.

**Table 1:** Adjusted Rand Index (ARI) scores of cell clustering based on the embedding learned by 10 different models from four gene+peak datasets (i.e., columns). For each dataset, we split the cells into 60% training and 40% test cells. The experiments were evaluated based on 500 random splits to record the mean and standard deviation of the performances of each method. When comparing between moETM and six SOTA methods for each dataset, the highest ARI scores is in bold and the second highest is in blue. The 10 models (i.e., rows) include six SOTA methods, our proposed moETM using PoE, moETM using MoE, and two single-omic ETM models trained on only RNA and ATAC, respectively.

| Metrics | Methods | Genes + Peaks |  |  |  |
| --- | --- | --- | --- | --- | --- |
|  |  | BMMC | MSLAC | MKC | MBC |
| ARI | SMILE | 0.719 (0.014) | 0.410 (0.012) | 0.409 (0.025) | 0.263 (0.025) |
|  | scMM | 0.693 (0.024) | 0.396 (0.014) | 0.392 (0.026) | 0.311 (0.019) |
|  | Cobolt | 0.678 (0.043) | 0.378 (0.014) | 0.373 (0.035) | 0.237 (0.048) |
|  | MultiVI | 0.703 (0.029) | 0.405 (0.020) | 0.382 (0.030) | 0.500 (0.031) |
|  | MOFA+ | 0.710 (0.012) | 0.421 (0.020) | 0.407 (0.041) | 0.244 (0.028) |
|  | Seurat V4 | 0.712 (0.008) | **0.504** (0.096) | 0.398 (0.025) | **0.507** (0.021) |
|  | moETM | **0.727** (0.007) | 0.469 (0.022) | **0.443** (0.025) | 0.423 (0.035) |
|  | moETM-average | 0.720 (0.004) | 0.460 (0.047) | 0.439 (0.047) | 0.410 (0.043) |
|  | moETM-rna | 0.720 (0.008) | 0.455 (0.053) | 0.433 (0.062) | 0.407 (0.054) |
|  | moETM-atac | 0.626 (0.009) | 0.252 (0.021) | 0.135 (0.008) | 0.188 (0.011) |

**Table 2:** Adjusted Rand Index (ARI) scores of cell clustering of 3 CITE-seq gene+protein datasets. Same as in **Table** 1, we split the cells into 60% training and 40% testing and evaluated each method based on how well their learned embedding cluster cells into known cell types.

| Metrics | Methods | Genes + Proteins |  |  |
| --- | --- | --- | --- | --- |
|  |  | BMMC | HWBC | HBIC |
| ARI | SMILE | 0.632 (0.020) | 0.397 (0.014) | 0.652 (0.030) |
|  | scMM | 0.655 (0.048) | 0.602 (0.039) | 0.670 (0.057) |
|  | Cobolt | 0.703 (0.036) | 0.572 (0.034) | 0.708 (0.043) |
|  | TotalVI | 0.744 (0.027) | 0.638 (0.051) | 0.733 (0.049) |
|  | MOFA+ | 0.713 (0.030) | 0.552 (0.038) | 0.687 (0.044) |
|  | Seurat V4 | 0.670 (0.009) | 0.553 (0.014) | 0.703 (0.025) |
|  | moETM | **0.760** (0.019) | **0.648** (0.023) | **0.752** (0.036) |
|  | moETM-average | 0.748 (0.023) | 0.626 (0.031) | 0.737 (0.049) |
|  | moETM-rna | 0.736 (0.030) | 0.617 (0.039) | 0.730 (0.040) |
|  | moETM-protein | 0.593 (0.015) | 0.523 (0.025) | 0.604 (0.042) |

For some topics, their top genes are both the members of the pathway and the cell-type biomarkers. For instance, topic 27 is enriched in the CD4+ T naïve cell. One of its top genes *CCR7* is involved in the elimination process of immature T cells. Additionally, topic 41 is enriched for the Transitional B cell. Its top pathways include B cell activation and adaptive immune process. Among its top genes, *TNFAIP3* is in the B cell activation-related pathway. One of its top peaks in chr14: 100207793 - 100208735 is upstream of the promoter of *YY1* (chr14: 100238298 - 100282788), which is a gene member in the B cell activation-related pathway [55].

Furthermore, we experimented a more specific gene set namely the immune signature gene set collections from MSigDB to investigate immune-related pathways implicated in the BMMC1 dataset (**Supplementary Fig. S4**). We identified several cell-type-specific topics that exhibit high scores for meaningful immune pathways. For instance, topic 23 is enriched in naïve CD20+ B cells. Two of its top 10 pathways are associated with naive B cells. One of its top genes namely *HLA-DPB1* is up-regulated in naïve B cells relative to the plasma cells [56]. One of the top peaks (chr12: 8886393 - 8887019) is upstream from *PHC1* (chr12:8913896 - 8941467), which is also involved in the pathway that genes are up-regulated in naive B cells relative to the plasma cells [56].

### 2.8. Multi-omic topics reveal the molecular basis of COVID-19 severity

As the CITE-seq technology interrogates the expression of surface proteins along with the full transcriptome, it is a promising platform to investigate the immune responses among patients infected by the SARS-CoV-2 virus (COVID-19). Using moETM, we sought to identify clinically relevant molecular signatures from a COVID-19 CITE-seq dataset (HBIC) [57]. The data consist of 781,123 cells from 130 COVID-19 patients with varying degrees of severity due to the viral infection. To establish model confidence, we first performed a quantitative analysis as above. The results showed that moETM could achieve either the highest or the second highest evaluation metrics both in bio-conservation and batch-removal cases (**Table** 2, **Supplementary Table S2**). In particular, moETM ranked first with an ARI value of 0.752 and TotalVI scored second with an ARI value of 0.733. Similarly, moETM and TotalVI attained the highest NMI scores of 0.779 and 0.762, respectively. Both methods also maintained their top performance in terms of batch-correction with TotalVI achieving the highest kBET of 0.197 while moETM coming in second with 0.153. Consistent to the above evaluation (**Table** S2), moETM obtained the best GC score of 0.950 whereas TotalVI achieved the second best of 0.934. Therefore, these quantitative results on the COVID-19 data further suggest that moETM strikes a good balance between biological conservation and batch effect correction in delivering competitive performance among all the SOTA methods.

Qualitatively, we investigated the top features and identified enriched cell types under each topic (**Supplementary Fig. S5a, b**). In particular, topic 42 is enriched for B cells. Among its top 5 genes (*SLC38A11*, *TCL1B*, *IL6*, *TCL1A*, *SYN3*), *IL6* and *TCL1A* are the known maker genes. Also, <u>3</u> out of its top 5 proteins (<u>CD19</u>, <u>CR1</u>, <u>CD22</u>, FCGR2A, BAFFR) are marker proteins for B cells. Topic 31 is associated with platelet. <u>Two</u> out of its top 5 genes (*LYVE1*, *RADIL*, *<u>VWF</u>* [58], *TRHDE*, *<u>PPBP</u>*) are marker genes, and <u>one</u> of its top 5 proteins (<u>ITGA2B</u>, KIR3DL1, ITGAX, SELP, FCGR2A) is a marker protein for platelet. Additionally, a previous study has suggested that SELP redistributes to the plasma membrane during platelet activation [59]. The enriched pathways based on GSEA are consistent to the cell-type specificity of those topics (**Supplementary Fig. S5c**). Taking topic 42 as an example, the enriched pathway is the gene set that is down-regulated in CD4 T cells compared with B cells [51]. Because of the shared embedding space, we also observed localization of the top genes and the top proteins for the selected topics via UMAP (**Supplementary Fig. S5d**).

We then leveraged the phenotype severity information among the patients to explore gene and protein signatures related to the COVID-19 phenotypes. Specifically, we utilized COVID metadata information to test whether a topic is significantly over-represented for the severity conditions. Here we considered each topic as a “meta-gene” and associated their up-regulation or down-regulation with the disease phenotypes (**Fig.** 7a, b). We delved into topics based on their distinctly enriched cell types, heatmap visualization (**Supplementary Fig. S5a**), and differential analysis by the phenotypes (e.g., COVID severity) (**Methods** 4.9). For example, we observed that topic 42 is not only enriched for B-cell but also up-regulated among patients with critical COVID status whereas topic 80 is significantly associated with the severe status. Moreover, topic 42 is associated with other demographic features such as age and mainly enriched in the senior group between 70 and 79 years of age (**Fig.** 7a).

**Figure 7:**
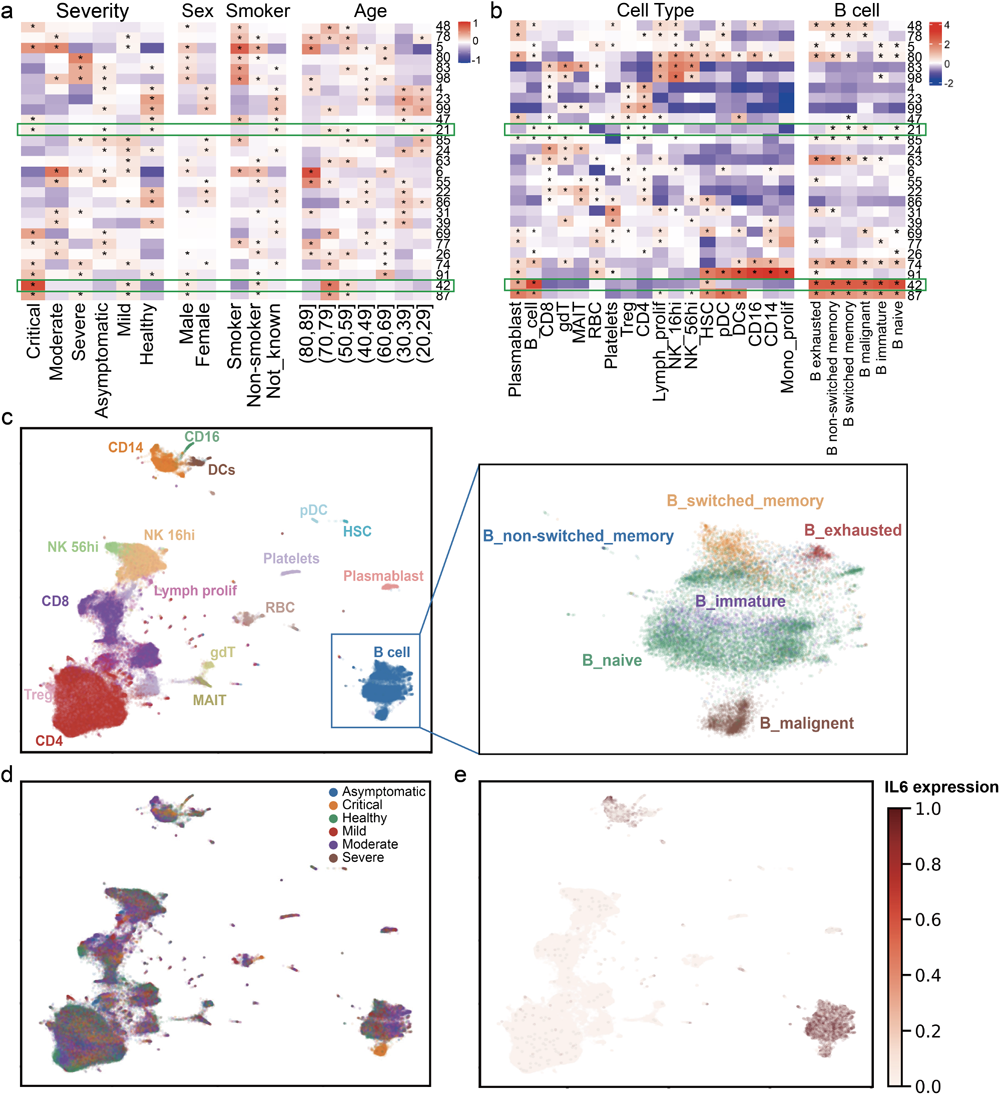
Topic association with the COVID-19 severity status. **a**. Differential analysis of severity states, sex, smoking history, and age. The color intensity values correspond to the differences of average topic scores between the positive cells and negative cells for each attribute (i.e., columns) and each topic (i.e., rows). Asterisks indicate Bonferroni-adjusted p-value *<* 0.001 based on one-sided t-test of up-regulated topics for each label. The results on the highlighted topic 21 and 42 were described in the main text. **b**. Differential analysis of topics across cell types. The heatmap on the left displays the topic associations with each of the 18 cell types, and the one on the right associates the same topics with 6 fine-grained B-cell subtypes. Similarly, asterisks indicate adjusted p-value < 0.001 for the t-test of up-regulated topics in each label. **c**. UMAP visualization of cell clustering. Colors indicate 18 cell types. The right panel shows a zoom-in version of the B-cell clustering with color indicating the 6 B-cell subypes. **d**. UMAP visualization with cells colored by source subjects’ severity states due to COVID-19 infection. **e**. Normalized gene expression of *IL6* among the cells on the same UMAP.

Given its disease relevance, we further investigated topic 42 to see whether it elucidates more granular cell types and to some extent whether their top gene/protein signatures can serve as putative biomarkers for COVID critical conditions. First of all, the moETM-inferred cell topic embeddings did not only cluster cells into their primary cell types but also sub-divided B cells into six sub-clusters of known sub-cell-types (**Fig.** 7c and zoom-in view). Intriguingly, aligning the COVID phenotypes with the B cell sub-types revealed that the critical COVID condition corresponded to B malignant cells (**Fig.** 7c, d). B-cell lymphomas start to develop when B lymphocytes, which are in charge of humoral immunity, start to proliferate beyond control. This proliferation turns B cells into malignant cells [60]. The previous study [61] suggested that individuals with certain cancers, such as lymphoma, may be more susceptible to getting severe illness from COVID-19. The top gene *IL6* in topic 42 was consistently expressed at a high level among B cells, including although not specifically in B malignant cells (**Fig.** 7e). *IL-6* levels were commonly reported in severely ill patients due to COVID-19 [62, 63]. As another example, topic 21 is also enriched in B malignant cells (**Fig.** 7b). One of its top proteins CD5 (**Supplementary Fig. S5b**) was shown to be highly expressed on malignant cells [64]. Moreover, the previous study [65] suggested that the proportion of CD5+ B cells was significantly reduced in COVID patients. Taken together, our results suggest that *IL-6* or CD5 may be a potential therapeutic target.

## 3 Discussion

Gene regulatory programs involve multi-faceted regulation and can not be understood via a single-omic approach alone. Single-cell multi-omic technologies open up the venues to interrogate several omics simultaneously in the same cells. As these technologies continue to evolve, computational methods are needed to account for the challenges in modeling the sparse, noisy, and heterogeneous nature of data that are being generated at a rapid pace [3]. In this study, we developed a unified interpretable deep learning model called moETM to integrate single-cell multi-omic data including transcriptome and chromatin accessibility or surface protein, which are the most common types of single-cell multi-omic data to date [4].

Our technical contributions are three-folds. First, via the product-of-experts, moETM effectively integrates multiple omics by projecting them onto a common topic mixture representation.

Second, the linear decoder enables the extraction of multi-omic signatures as the top features under each latent topic, which directly reveal marker genes and phenotype markers under topics that are aligned with cell types or phenotype conditions. Third, by efficiently correcting batch effects via a dedicated linear intercept matrix in the decoder, we can integrate multi-omic data from multiple studies, subjects, or technologies, which allows us to exploit the vast amount of multi-omic data in order to obtain biologically diverse and coherent multi-omic topics. Notably, while the last two contributions are inherited from our earlier single-cell transcriptome model namely scETM [15], we consider the success of incorporating them into the multi-omic modeling problem as a substantial departure from the existing method.

To demonstrate the utility of moETM, we benchmarked it with 6 existing state-of-the-art computational methods on 7 published datasets including 4 gene+peak datasets and 3 gene+protein datasets (**Table** 1, 2). Across all datasets, moETM achieved competitive performance based on 4 common evaluation metrics including the bio-conservation of recovering known cell types (i.e., ARI and NMI) and batch-removal evaluation metrics (i.e., kBET and GC). We also confirmed the advantage of using multiple modalities compared with single modality in terms of cell clustering (**Table** 1, 2, **Supplementary Table S1, S2**).

As the vast majority of the single-cell data are still single-omic (e.g., scRNA-seq, scATAC-seq, etc), there are tremendous benefits of imputing one omic from another omic. Because of its joint modeling capabilities, the trained moETM can accomplish this cross-omic imputation task. The imputation can go from a high-dimensional omic to a low-dimensional omic and vice versa. The latter imputation direction is more demanding on the decoder because it needs to “remember” the high-dimensional manifold in its parameter space when decoding the lower-dimensional feature space. In our applications, this involves imputing gene expression (*∼*20K) from ATAC peaks (*∼*100K) and imputing surface protein abundance (*∼*140) from gene expression (*∼*20K) and vice versa. In both imputation directions, moETM achieved a higher correlation than scMM and BABEL. Although more challenging, moETM also achieved a reasonable performance when imputing high-dimension from low-dimension.

We also explored the moETM-learned cell-type-specific topics in terms of their top omic features and enriched pathways in the light of the supporting evidence from the literature. For example, in the BMMC2 (gene+protein) dataset, *CR2* is a top gene signature identified by high topic scores in a B-cell-specific-topic topic 40, a marker gene for B cell in the CellMarker database (**Fig.** 5b, c), and a member of an enriched pathway (genes that are down-regulated in CD4 T cells compared with B cells [51]) for the topic. By binding to C3d, *CR2* can lower the threshold for B cell activation in an adaptive immune response [66]. Similarly, in the BMMC2 (gene+protein) dataset, protein CD19 is included in the topic enriched pathway (genes that are down-regulated in CD4 T cells compared with B cells [51]). It is ranked sixth in the B-cell-specific topic 40 and is crucial in determining intrinsic B cell signaling thresholds. Along with other molecules, CD19 functions as the dominant signaling element of a multimolecular complex on the surface of mature B cells [67]. In the BMMC1 (gene+peak) dataset, a top gene feature *IL4I1* in a T-cell-specific topic 3 is a target gene of the enriched motif FLI1 (**Fig.** 6d), which is determined by the top 100 peaks under that topic. *IL4I1* could inhibit human CD4+ and CD8+ T lymphocyte proliferation *in-vitro* [68]. Furthermore, T cells have a high-level expression of FLI1 and the expression decreases after T cell activation [69].

In a more focused study, we analyzed the COVID-19 CITE-seq dataset (gene+protein) and linked moETM-learned immune-specific topics with patient severity conditions due to the infection. Our topic analysis revealed not only immune marker genes but also cell types that are associated with COVID phenotype conditions. In particular, we found that the patients with critical status exhibited high topic probabilities for the B malignant cells. Furthermore, one of the B malignant cell marker genes *IL6* is differentially expressed among these patients compared to patients with mild and no symptoms ([62, 63]).

There are several challenges that are not addressed in moETM [4]. For instance, moETM has the capacity to integrate across multiple batches and modalities but it requires the training data to have all omics measured in the same cells. Given that the transcriptome is shared between the gene+peak and gene+protein data, it is possible to integrate all 3 omics in a mosaic data integration regime while taking into account the data heterogeneity. A more challenging task is to integrate multimodal data without anchored features or cells, which is commonly known as the diagonal integration [4]. Some recent approaches made use of graph representation learning to integrate multi-omic single-cell data at the expense of computational complexity and interpretability [70–72]. Furthermore, given the motif enrichments in our analysis, another natural extension of moETM is to model the sequence information at the upstream of the model training using language models such as the Bidirectional encoder representations from transformer (BERT) model [73]. Indeed, at the decreasing computational cost, we started to see interesting applications of BERT in the related fields including single-cell data modeling [74], genome language understanding [75], and sequence-based gene expression prediction [76].

## 4 Methods

### 4.1. moETM data generative process

The molecular activities in each cell *n* can be measured with *M* omics, such as gene expression from transcriptome, surface protein expression, and the open chromatin regions manifested as peaks. For the ease of the following descriptions, we define the entities of genes, proteins and peaks as “features”. Profiling those omics in the cell leads to *M* count vectors, each of which has a dimension *V* ^(^*^m^*^)^ as the number of unique features in omic *m*. Adapting the text-mining analogy, we consider each cell as a “document” written in *M* languages or modalities (i.e., transcriptome, proteome, chromatin accessibility); each feature from the *m^th^* omic is considered as a “word” from the *m^th^* vocabulary; each sequencing read is a “token” in the document; the abundance of the reads mapped to the same feature is the “word count” in the document.

The multi-modal document of a cell *n* can be summarized into a mixture of *K* latent topics ***θ****_n_*, which are presumably implicated in each modality (**Fig.** 1a). Inference of these topic mixtures for each cell is accomplished by modeling the distribution of the multi-omic count data *{***x**^(^*^m^*^)^*}^M^* from the topic mixture for the cell and learning the global topic embedding over the *M* modalities. The latter are shared among all cells and expressed as *M* matrices, where a row vector ***ϕ***^(^*^m^*^)^ *∈* R*^V^* ^(^*^m^*^)^ denotes the *k*-th topic from the *m*-th modality.

To increase information sharing across the omics and the model expressiveness, we further decompose each omic-specific topic embedding matrix **Φ**^(^*^m^*^)^ into the topic embedding ***α*** *∈* R*^K×L^* and feature embedding ***ρ***^(^*^m^*^)^ *∈* R*^L×V^* ^(^*^m^*^)^, where *L* denotes the size of the embedding space. The expected values for the count data for each omic is proportional to the dot product of the cell embedding, topic embedding matrix, and feature embedding matrix: **x**^(^*^m^*^)^ ***θ****_n_*

Formally, we formulate the data generative process as follows. For each cell indexed by *n ∈ {*1*, . . ., N}*, draw a 1 *× K* topic proportion ***θ****_n_* from logistic normal distribution ***θ****_n_ ∼ LN* (**0**, **I**):

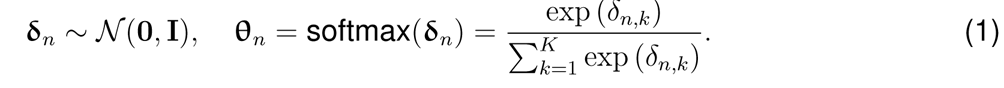

For each read *i*^(^*^m^*^)^ *∈ {*1*, . . ., D*^(^*^m^*^)^*}* from the *m^th^* modality *w*^(^*^m^*^)^, draw a feature index *v*^(^*^m^*^)^ (e.g., *n _n,i_*(*m*) the particular transcript or open chromatin region the read was sequenced) from a categorical distribution Cat(**r**^(^*^m^*^)^):

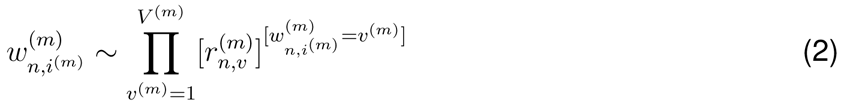

where *D*^(^*^m^*^)^ denotes the total number of reads. The expected rate *r*^(^*^m^*^)^ of observing feature *v*^(^*^m^*^)^ in cell *n* is parameterized as:

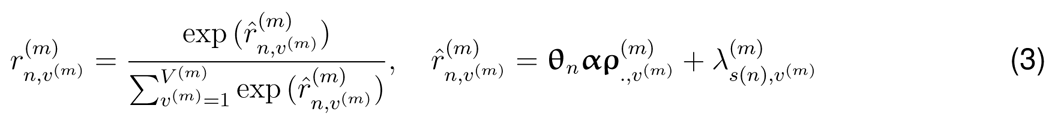

where *∈* R*^L×^*^1^ denotes embedding of feature *v*^(^*^m^*^)^, *λ*^(^*^m^*^)^ is the batch-dependent and feature-specific scalar effect, where *s*(*n*) indicates the batch index for the *n^th^* cell. Notably, the softmax function normalizes the expected observation rates over all features separately within each modality to account for different modality size (e.g., there are more peaks than genes, and more transcripts than surface proteins). Another reason for the normalization is to capture feature sparsity (i.e., only a small fraction of features from each modality is non-zero). This is analogous to text mining, where only a small fraction of the unique words are draw from the entire vocabulary for any given document.

The likelihood for cell *n* can be expressed as multinomial distribution:

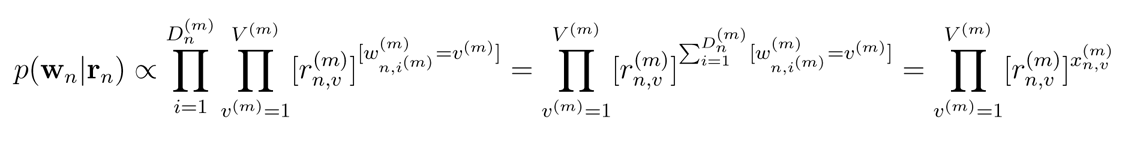

where 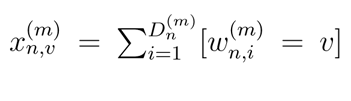denotes the read count for feature *v* for cell *n* in the *m^th^* modality. As a result, we can more conveniently express the data likelihood in terms of *the read count*:

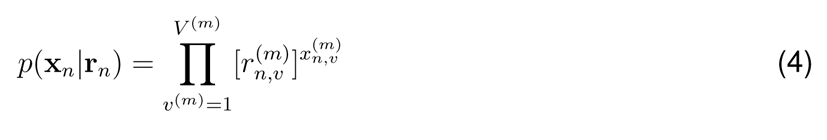

### 4.2. moETM model inference

For the ease of inference, we consider the cell topic embedding ***δ****_n_* (before softmax normalization) for cells *n ∈ {*1*, . . . N}* as the latent variables and all the cells are independent. The rest of the parameters including topic embedding ***α***, feature embedding *{****ρ***^(^*^m^*^)^*}^M^*, and batch-effect parameter *{****λ***^(^*^m^*^)^*}^M^* are treated as point estimates and learned by the model. Let’s denote Θ^^^ = *{****δ****_n_, **α**, {****ρ***^(^*^m^*^)^*}^M^* *, {****λ***^(^*^m^*^)^*}^M^*)*}*. A principled way to learn those parameters is to maximize the log marginal likelihood:

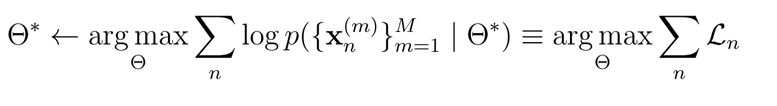

However, this integral is not tractable. Instead, we took a variational inference approach to optimize the model parameters by maximizing an evidence lower bound (ELBO) of the marginal log likelihood with a proposed variational posterior *q*(***δ****_n_*) as a surrogate to the true posterior of the cell topic embedding *p*(***δ*** *| {***x**^(^*^m^*^)^*}^M^*):

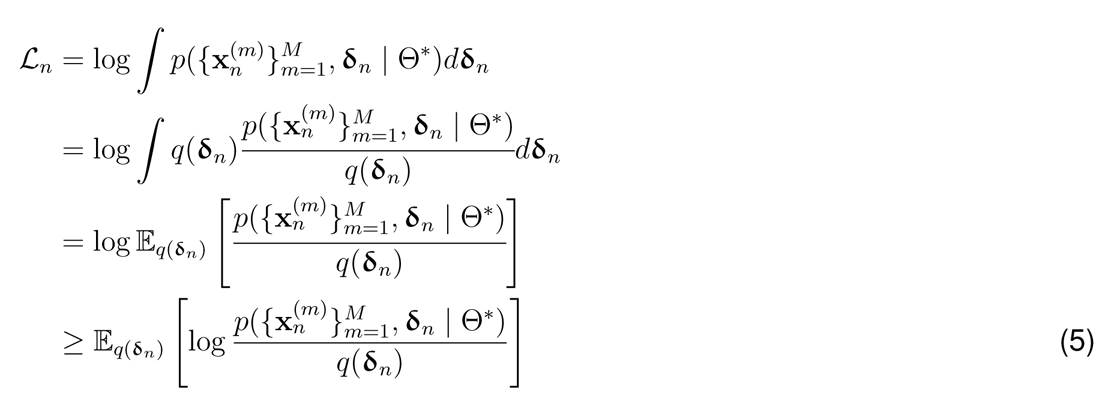

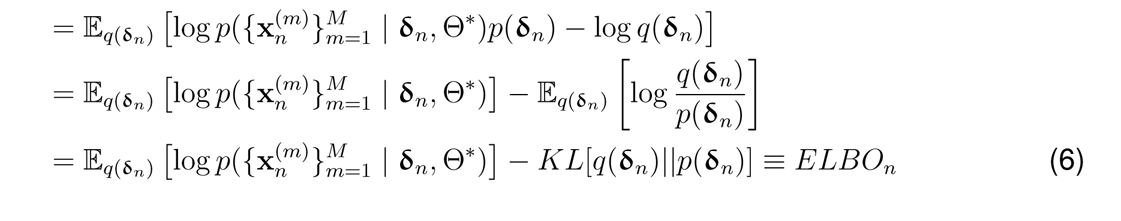

where Eq (5) follows the Jensen’s inequality [77] and KL denotes the Kullback-Leibler (KL) divergence between the proposed distribution and the prior (i.e., standard Gaussian with zero mean and identity variance), acting as a regularization when maximizing the data likelihood.

We defined the proposed distribution *q*(***δ****_n_*) as a product of Gaussians (PoGs):

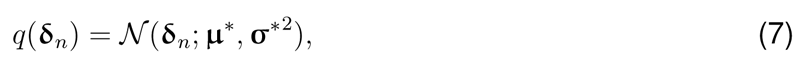

The mean ***µ*** and standard deviation ***σ*** of the joint Gaussian is computed as:

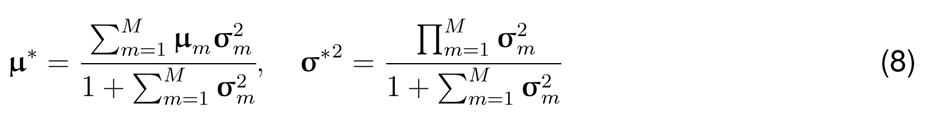

where ***µ****_m_* and ***σ***^2^ are the mean and variance of the Gaussian latent embedding for the individual modalities, respectively. Those are output from the encoder neural network (NNET):

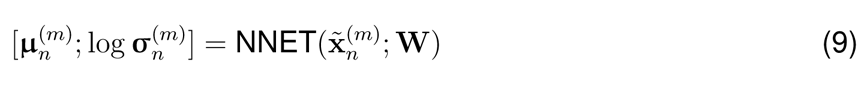

where **x**”^(^*^m^*^)^ is the normalized counts for each feature as the raw count of the feature divided by the total counts of *m^th^* modality in cell *n*, and **W** is the parameters for a two-layer feed-forward neural network.

We approximate the above ELBO in Eq (6) by sampling from the proposed joint Gaussian distribution using the reparameterization trick [13]:

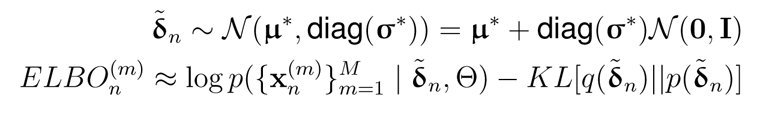

where the KL divergence has closed form:

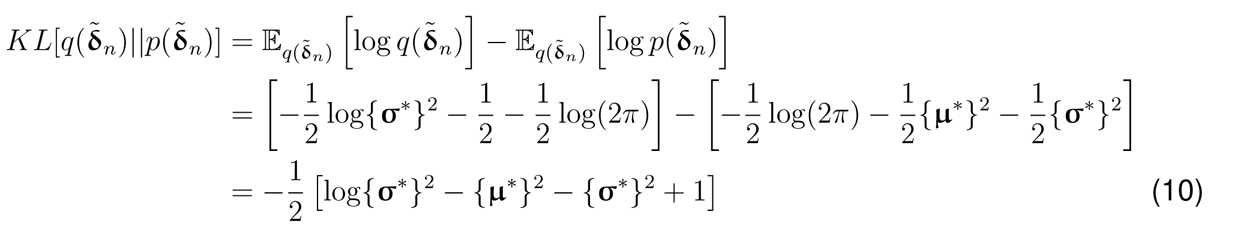

Together, with the Multinomial likelihood defined in (4) and KL divergence in (10), we can express the ELBO in its approximate closed-form using the sampled latent variable:

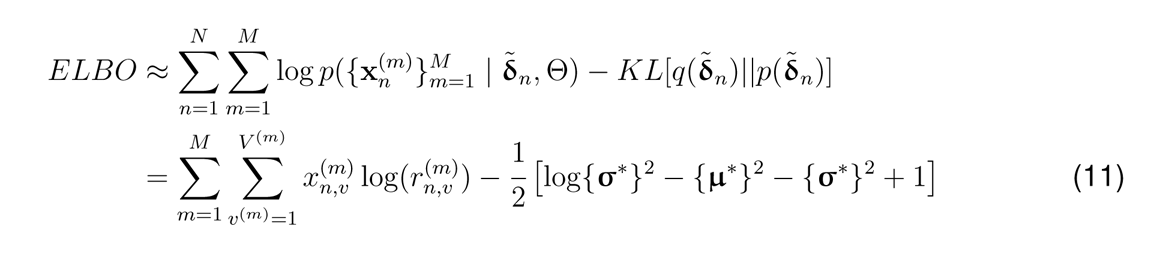

where *r*^(^*^m^*^)^ is defined in (3). The model parameters including the encoder weight **W** and the de-coder weights Θ = *{****α****, **ρ**}* are optimized by maximizing the above ELBO via backpropagation:

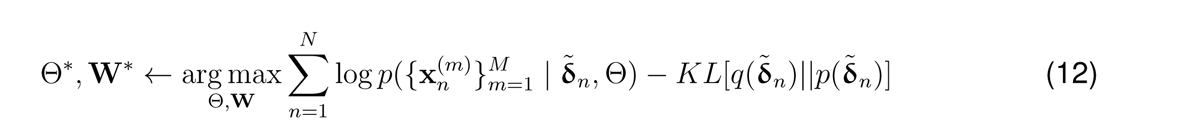

### 4.3. Single-cell multi-omic datasets and preprocessing

There were 7 public datasets included in this study for performance evaluation and model comparison. All 7 datasets are from publicly available repositories. Among them, 4 datasets provide joint profiling of gene expression and open chromatin regions (denoted as “gene+peak” data):

1. Multiome bone marrow mononuclear cells (BMMC1) dataset from the 2021 NeurIPS challenge consisting of 42,492 cells with 22 cell types from 10 donors across 4 sites [78],
2. SHARE-seq mouse skin late anagen (MSLAC) dataset containing 34,774 cells with 1 batch and 23 cell types [24],
3. sci-CAR mouse kidney cells (MKC) dataset from cell samples with 1 batch and 14 cell types [79],
4. SHARE-seq mouse brain cells (MBC) dataset containing 3,293 cells with 1 batch and 19 cell types [24].

For the BMMC1 dataset, we take into account two different batch types: one treats a subject (eg. site1 + donor1 as a subject s1d1, site1 + donor2 as a subject s1d2, etc) as a batch (s1d1, s1d2, s1d3, s2d1, s2d4, s2d5, s3d3, s3d6, s3d7, s3d10, s4d1, s4d8, s4d9, 13 batches in total), while the other treats a site (site1 as batch1, site2 as batch2) as a batch (4 batches in total).

For the CITE-seq data measuring transcriptome and surface protein in the same cell, 3 datasets were used in this study:

1. Bone marrow mononuclear cells (BMMC2) dataset from the 2021 NeurIPS challenge from 9 donors and 4 sites [78],
2. Human White Blood Cell (HWBC) dataset containing 211,000 human peripheral blood mononuclear cells [12],
3. Human Blood Immune Cell (HBIC) dataset [57] measuring 647366 peripheral blood mononuclear cells from both COVID patients and healthy patients.

Similarly, for the BMMC2 dataset, we consider two different batch types: one treats donors as batches (12 batches in total), while the other treats sites as batches (4 batches in total).

All datasets were processed into the format of samples-by-features matrices. For gene+peak datasets, the read count for each gene and peak were first normalized per cell by total counts within the same omic using *scanpy.pp.normalize_total* function in the *scanpy* [80], then log1p transformation was applied. After that, *scanpy.pp.highly_variable_genes* was used to select highly variable genes or peaks.

For the joint profiling of transcriptome and surface protein data (denoted as gene+protein), we used all surface proteins measured by the scADT-seq assay since the number of proteins is much smaller compared with the number of genes or peaks and all of them are highly informative of immune cell functions. The same normalization as in the gene+peak data was performed on the gene+protein data.

### 4.4. Cross-omic imputation

The trained moETM can impute one omic from another omic. Suppose we have two omics namely omic A and omic B. For the training data where both omics are observed, moETM learns a shared topic embedding ***α*** and omic-specific feature embedding ***ρ***^(^*^A^*^)^ and ***ρ***^(^*^B^*^)^. For the testing data, suppose without loss of generality that only omic B is observed. To impute omic A, moETM uses the encoder for modality B to generate the topic mixture, which is then input to the decoder for omic A to complete the imputation (**Fig.** 1c).

We evaluated the imputation accuracy using the BMMC1 (gene+peak) and BMMC2 (gene+protein) datasets based on (1) 60/40 random split of training and testing data with 500 repeats to get standard deviation estimate; (2) training on all batches except for one batch and testing on the held-out batch (leave-one-batch); (3) training on all cell types except for one cell type and testing on the held-out cells of that cell type (leave-one-cell-type).

### 4.5. Evaluation metrics

The batch effects correction and biological variance conservation categories were used to assess the efficacy of the integration across multiple modalities. To quantify bio-conservation, we used the Adjusted Rand Index (ARI) and Normalized Mutual Information (NMI), and to measure batch effect removal, we used k-nearest-neighbor batch-effect test (kBET) and Graph Connectivity (GC). Specifically, ARI calculates the degree of similarity between two clusterings and adjusts for the possibility that objects can randomly form the same clusters. NMI normalizes the mutual information to a scale of 0 to 1. While NMI excels in unbalanced clustering or small clusters, ARI is better suited to clusters of similar size [81]. kBET performs hypothesis testing on whether batch labels are distributed differently across cells based on Pearson’s *χ*^2^ test [19]. GC measures whether cells of the same type from different batches are close to one another by computing a K nearest-neighbour graph based on the distance between cells in the embedding space [20].

### 4.6. Linking genes to open chromatin regions

We sought to investigate the relation between the top peaks and top genes under the same moETM topic (i.e., ***ϕ***^(^*^m^*^)^ = ***α****_k_**ρ***^(^*^m^*^)^ for topic *k* and *m ∈ {gene, peak}*). To assess the *in-cis* relation, we measured the genomic distances between genes and peaks and designated genes that were near peaks as peak-neighboring-genes if they are within 150K base pairs (bp) distance.

Specifically, we first obtained a genes-by-topics matrix ***ϕ***^(gene)^ = ***αρ***^(gene)^ and a peaks-by-topics matrix ***ϕ***^(peak)^ = ***αρ***^(peak)^.

To transform ***ϕ***^(peak)^ into a peak_to_genes-by-topics matrix ***ϕ***^(peaks_to_genes)^, we first derived a binary peaks-to-genes mapping matrix **H** with the entries *h_p,g_* = 1 if the corresponding pair of peak *p* and gene *g* are within 150K bp genomic distance and are positively correlated and 0 otherwise.

In detail, we computed the Pearson correlation between gene *g* and peak *p* in terms of their topic scores:

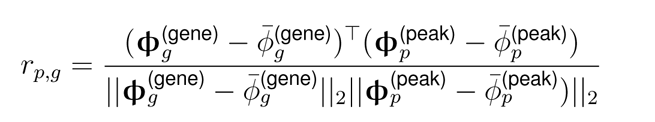

The genome distance between peaks and genes was based on the latest genome build (i.e., hg38 for human) and obtained via the *GenomicRanges* [82] package in R.

### 4.7. Pathway enrichment analysis

For each moETM topic, we performed Gene Set Enrichment Analysis (GSEA) [37] to associate the topic with known pathways or gene sets. In particular, we used each topic to query two gene sets from Molecular signatures database (MSigDB), which are the 5219 Immunologic signature gene sets (C7) and the 7763 Gene Ontology Biological Processes (BP) (C5-BP) terms. For each topic, we ran *GSEAPreranked* on a ranked list of genes based on their corresponding topic scores against every gene set from C7 or C5-BP, and calculated the enrichment score (ES) for over- or under-representation. The statistical significance of the ES was computed based on 1000 permutation test. The gene sets with Benjamini–Hochberg (BH) corrected p-values lower than 0.05 were deemed significant. Similarly, for the scATAC-seq data, the peaks-by-topics matrix was first converted into a peaks_to_genes-by-topics matrix and then provide as input to GSEA pipeline.

### 4.8. Motif enrichment analysis of top peaks from moETM-learned topics

To detect sequence-based regulatory elements for the cell-type-specific topics, we performed motif enrichment analysis using the top 100 peaks that exhibit the highest topic scores under each topic. The 100 sequences corresponding to those top 100 peaks under each topic were extracted from Ensembl database and provided as input to the Simple Enrichment Analysis (SEA) pipeline [83] from the MEME suite [84]. SEA utilizes the STREME motif discovery algorithm [85] to identify known motifs that are enriched in input sequences. For our purpose, we used the HOmo sapiens COmprehensive MOdel COllection (HOCOMOCO) Human (v11) and HOCOMOCO Mouse (v11) motif database [86]. Motifs with Fisher’s exact test p-values lower than 0.05 were selected as the enriched motifs.

### 4.9. Differential analysis to detect condition-specific topics

We sought to detect moETM-topics that exhibit significantly higher scores for the conditions of interest such as cell types or phenotypes. Notably, while the cell types were at the single-cell level, the phenotypes were at the subject level (e.g., COVID-19 severity state). The latter means that the cells from the same subject were assigned the same phenotype label. For each dataset, we first split the cells into positive and negative groups, corresponding to the presence and absence of the target condition, respectively. For each topic, we assessed the statistical significance of the topic score increase for the positive group relative to the negative group based on one-sided student t-test. The topics with a Bonferroni-adjusted p-value smaller than 0.001 were considered significant with the label.

### 4.10. Incorporating pathway-informed gene embeddings

In the linear decoder, we reconstruct the cells-by-features matrix by the dot product of the 3 matrices, namely cells-by-topics, topics-by-embedding, and embedding-by-features. By default, the last feature embedding matrix consist of learnable parameters. However, we can instill prior pathway information during the training of moETM by fixing the features embedding to a known gene set. As a result, the topics-by-embedding and embedding-by-features matrices change to topics-by-gene_sets and gene_sets-by-features with only the topics-by-gene_sets as the learn-able parameters. This allows us to directly map each topic to each gene set, which may further improve the model interpretability especially if the chosen gene sets were highly relevant to the data. Given that several single-cell multi-omic datasets used in this study were derived from the blood, we utilized the Immunologic signature gene sets collection (C7) from the MSigDB database. Gene sets with fewer than five or more than 1000 genes were filtered out. We then converted the gene set information into a binary gene_sets-by-genes matrix with 0 and 1 indicating the absence and presence of the genes (columns) in the corresponding gene set (rows), respectively. We focused on the gene+peak case by fixing the gene embedding to the gene set while learning the peak embedding as in the default setting. We did not experiment this approach on the gene+protein case, for which the topics learned by the default moETM are sufficiently easy to interpret.

## Code Availability

The moETM code is available at https://github.com/manqizhou/moETM.

## Acknowledgement

F.W. would like to acknowledge the support from NIH R01AG076448, R01AG076234, RF1AG072449, NSF 1750326 and 2212175. Y.L. is supported by NSERC Alliance Catalyst ALLRP 576153-22, NSERC Discovery Grant DGECR-2019-00253 and CIHR Canada Research Chair (Tier 2) in Machine Learning for Genomics and Healthcare.

## Author contributions

M.Z. and H.Z. designed and performed the experiments under supervision of F.W. and Y.L. All authors read and approved the final manuscript.

## Supplementary Information

### S1 Supplementary Tables

**Table S1:**
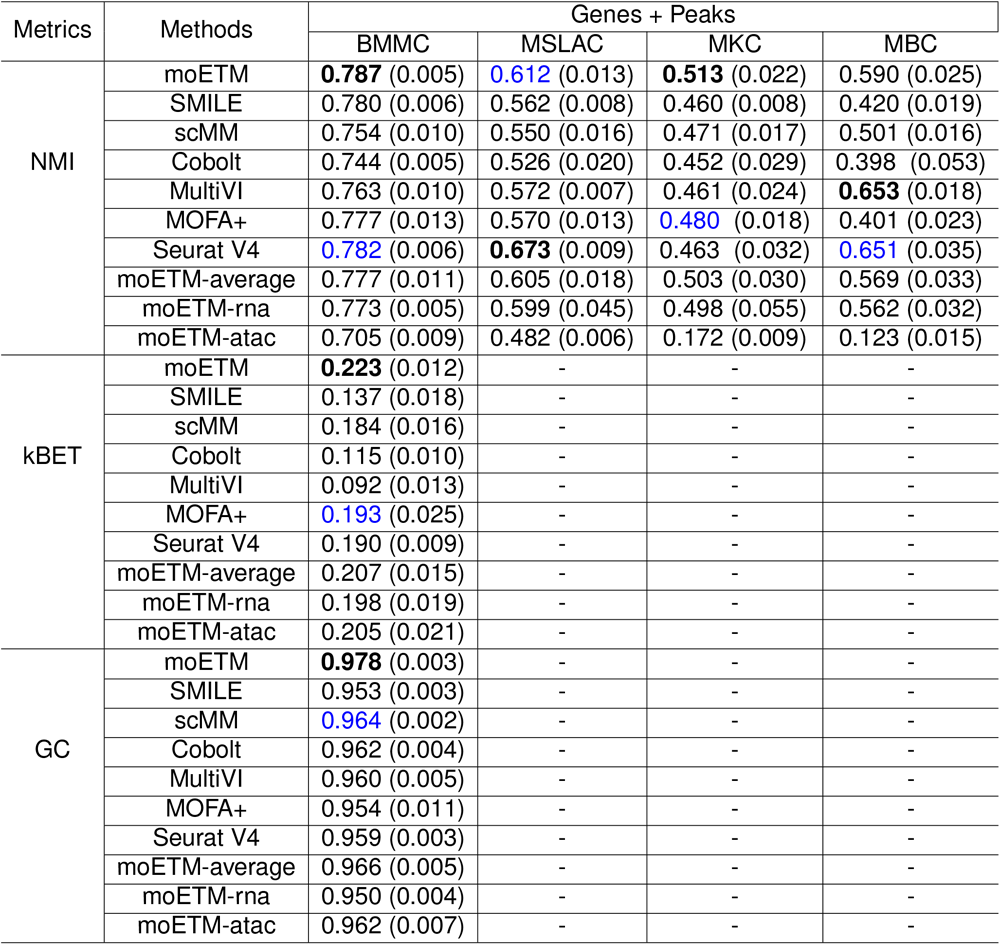
Evaluation of cell clustering by NMI, kBET, and GC on 4 genes+peaks single-cell multi-omic datasets. The experiments were the same as described in **Table** 1. Only the BMMC data have multiple batches (13 batches) and therefore evaluated by the kBET and GC scores. The other 3 datasets were evaluated by NMI and ARI only based on their cell-type labels (**Table** 1).

**Table S2:** Evaluation of cell clustering of 3 CITE-seq gene+protein datasets based on NMI, kBET, and GC. The experiments were the same as described in **Table** 2.

| Metrics | Methods | Genes + Proteins |  |  |
| --- | --- | --- | --- | --- |
|  |  | BMMC | HWBC | HBIC |
| NMI | moETM | **0.786** (0.006) | **0.791** (0.013) | **0.779** (0.023) |
|  | SMILE | 0.712 (0.007) | 0.732 (0.005) | 0.702 (0.017) |
|  | scMM | 0.708 (0.012) | 0.742 (0.018) | 0.712 (0.034) |
|  | Cobolt | 0.731 (0.011) | 0.730 (0.027) | 0.739 (0.030) |
|  | TotalVI | **0.770** (0.005) | **0.784** (0.010) | **0.762** (0.019) |
|  | MOFA+ | 0.750 (0.024) | 0.701 (0.027) | 0.715 (0.029) |
|  | Seurat V4 | 0.714 (0.006) | 0.714 (0.040) | 0.726 (0.012) |
|  | moETM-average | 0.770 (0.011) | 0.769 (0.018) | 0.753 (0.029) |
|  | moETM-rna | 0.755 (0.004) | 0.759 (0.010) | 0.748 (0.019) |
|  | moETM-protein | 0.625 (0.013) | 0.620 (0.019) | 0.608 (0.027) |
| kBET | moETM | **0.105** (0.002) | **0.304** (0.012) | **0.153** (0.020) |
|  | SMILE | 0.058 (0.001) | 0.032 (0.007) | 0.073 (0.017) |
|  | scMM | 0.080 (0.004) | 0.110 (0.007) | 0.092 (0.011) |
|  | Cobolt | 0.036 (0.002) | 0.095 (0.005) | 0.038 (0.019) |
|  | TotalVI | **0.156** (0.009) | **0.349** (0.002) | **0.197** (0.004) |
|  | MOFA+ | 0.078 (0.013) | 0.213 (0.010) | 0.063 (0.014) |
|  | Seurat V4 | 0.062 (0.003) | 0.133 (0.010) | 0.107 (0.015) |
|  | moETM-average | 0.094 (0.005) | 0.290 (0.016) | 0.140 (0.023) |
|  | moETM-rna | 0.082 (0.002) | 0.266 (0.010) | 0.130 (0.015) |
|  | moETM-protein | 0.090 (0.006) | 0.280 (0.015) | 0.135 (0.023) |
| GC | moETM | **0.936** (0.007) | **0.968** (0.004) | **0.950** (0.005) |
|  | SMILE | 0.901 (0.011) | 0.912 (0.016) | 0.907 (0.006) |
|  | scMM | 0.897 (0.009) | 0.937 (0.016) | 0.906 (0.008) |
|  | Cobolt | 0.880 (0.011) | 0.930 (0.019) | 0.890 (0.010) |
|  | TotalVI | **0.918** (0.005) | **0.951** (0.006) | **0.934** (0.007) |
|  | MOFA+ | 0.906 (0.022) | 0.940 (0.022) | 0.895 (0.015) |
|  | Seurat V4 | 0.911 (0.004) | 0.930 (0.002) | 0.923 (0.003) |
|  | moETM-average | 0.920 (0.009) | 0.949 (0.006) | 0.936 (0.008) |
|  | moETM-rna | 0.911 (0.010) | 0.932 (0.007) | 0.928 (0.009) |
|  | moETM-atac | 0.916 (0.011) | 0.939 (0.007) | 0.930 (0.015) |

**Table S3:**
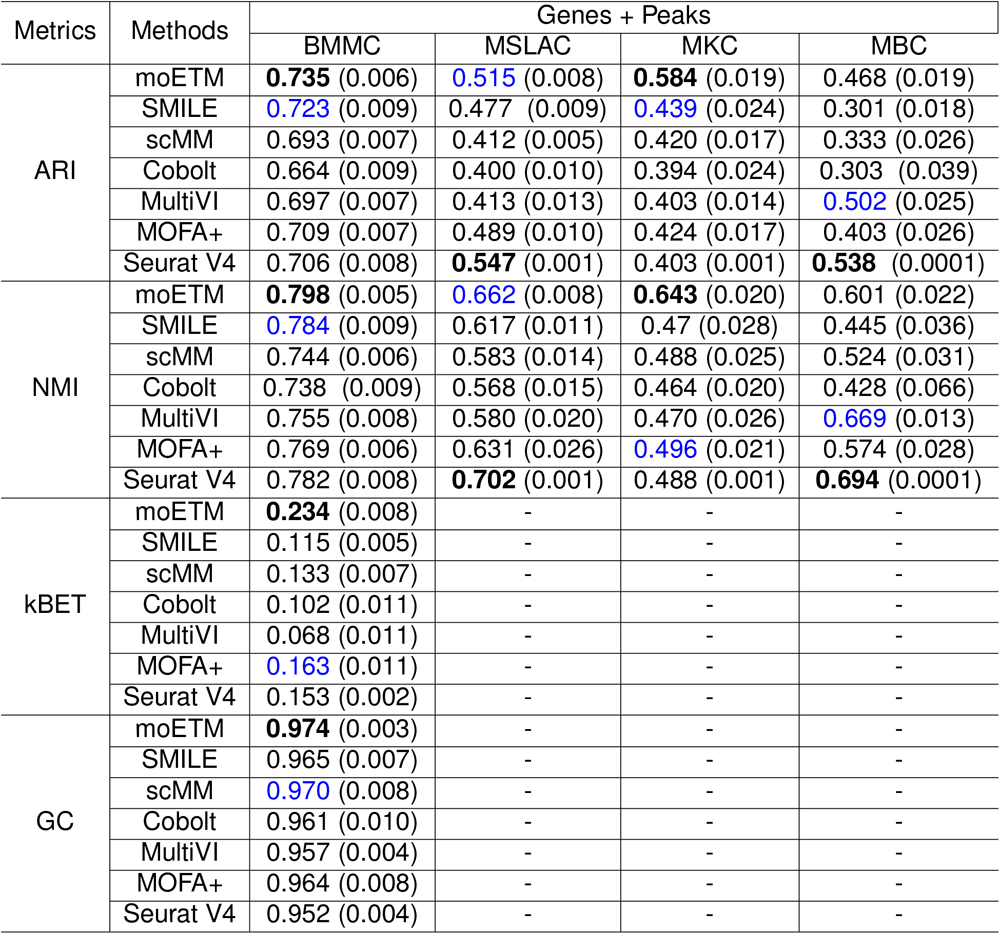
Evaluation of embedding-based cell clustering of the 4 gene+peak datasets. Different from the one listed in **Table** 1 and **Supplementary Table** S1, we trained and tested each model on all of the cells from each dataset. Because the cell type labels were not used in training any of the model, this is still an unbiased evaluation.

**Table S4:** Evaluation of embedding-based clustering by leaving-one-**subject**-out (Section 4.3). Each method was trained on *B −* 1 subjects and tested on the held-out subject. The highest and second highest score per dataset were highlighted in bold and blue, respectively.

| Metrics | Methods | Genes + Peaks | Genes + Proteins |
| --- | --- | --- | --- |
|  |  | BMMC1 | BMMC2 |
| ARI | moETM | **0.779** (0.071) | **0.776** (0.071) |
|  | SMILE | 0.766 (0.082) | 0.703 (0.099) |
|  | scMM | 0.743 (0.120) | 0.737 (0.084) |
|  | Cobolt | 0.732 (0.105) | 0.726 (0.013) |
|  | MultiVI/TotalVI | 0.739 (0.095) | 0.740 (0.107) |
|  | MOFA+ | 0.759 (0.098) | 0.713 (0.034) |
|  | Seurat V4 | 0.730 (0.095) | 0.686 (0.120) |
| NMI | moETM | **0.819** (0.037) | **0.823** (0.029) |
|  | SMILE | 0.808 (0.036) | 0.803 (0.030) |
|  | scMM | 0.780 (0.056) | 0.804 (0.026) |
|  | Cobolt | 0.750 (0.150) | 0.799 (0.034) |
|  | MultiVI/TotalVI | 0.774 (0.043) | 0.810 (0.038) |
|  | MOFA+ | 0.783 (0.042) | 0.758 (0.027) |
|  | Seurat V4 | 0.782 (0.050) | 0.744 (0.023) |

**Table S5:** Evaluation of embedding-based clustering by leaving-one-**site**-out (Section 4.3). Each method was trained on *B −* 1 sites and tested on the held-out site. The highest and second highest score per dataset were highlighted in bold and blue, respectively.

| Metrics | Methods | Genes + Peaks | Genes + Proteins |
| --- | --- | --- | --- |
|  |  | BMMC1 | BMMC2 |
| ARI | moETM | **0.735** (0.084) | **0.772** (0.053) |
|  | SMILE | 0.714 (0.095) | 0.686 (0.075) |
|  | scMM | 0.700 (0.106) | 0.665 (0.066) |
|  | Cobolt | 0.693 (0.068) | 0.653 (0.057) |
|  | MultiVI/TotalVI | 0.704 (0.086) | 0.694 (0.085) |
|  | MOFA+ | 0.716 (0.078) | 0.668 (0.073) |
|  | Seurat V4 | 0.693 (0.097) | 0.666 (0.149) |
| NMI | moETM | **0.796** (0.036) | **0.810** (0.033) |
|  | SMILE | 0.780 (0.041) | 0.798 (0.040) |
|  | scMM | 0.760 (0.033) | 0.771 (0.022) |
|  | Cobolt | 0.768 (0.062) | 0.759 (0.043) |
|  | MultiVI/TotalVI | 0.762 (0.038) | 0.787 (0.050) |
|  | MOFA+ | 0.780 (0.042) | 0.768 (0.041) |
|  | Seurat V4 | 0.756 (0.038) | 0.737 (0.010) |

**Table S6:** Imputing surface protein expression from gene expression using the BMMC2 dataset. The best score per evaluation metric is in bold.

| Methods | random split |  | leave-one-batch |  | leave-one-cell-type |  |
| --- | --- | --- | --- | --- | --- | --- |
|  | Pearson | Spearman | Pearson | Spearman | Pearson | Spearman |
| moETM | **0.95** (0.02) | **0.94** (0.02) | **0.92** (0.02) | **0.90** (0.03) | **0.88** (0.04) | **0.85** (0.03) |
| BABEL | 0.94 (0.03) | 0.92 (0.03) | 0.90 (0.02) | 0.87 (0.03) | 0.85 (0.03) | 0.81 (0.03) |
| scMM | 0.94 (0.03) | 0.91 (0.04) | 0.91 (0.03) | 0.89 (0.03) | 0.83 (0.04) | 0.78 (0.05) |

**Table S7:** Imputing gene expression from chromatin accessibility using the BMMC1 dataset. The best score per evaluation metric is in bold.

| Methods | random split |  | leave-one-batch |  | leave-one-cell-type |  |
| --- | --- | --- | --- | --- | --- | --- |
|  | Pearson | Spearman | Pearson | Spearman | Pearson | Spearman |
| moETM | **0.69** (0.02) | **0.37** (0.03) | **0.65** (0.04) | **0.35** (0.04) | **0.58** (0.05) | **0.32** (0.03) |
| BABEL | 0.65 (0.03) | 0.34 (0.03) | 0.6 (0.03) | 0.33 (0.03) | 0.55 (0.05) | 0.30(0.02) |
| scMM | 0.63 (0.03) | 0.33 (0.04) | 0.61 (0.04) | 0.33 (0.03) | 0.54 (0.05) | 0.28 (0.04) |

**Table S8:** Imputing chromatin accessibility from gene expression using the BMMC1 dataset. The best score per evaluation metric is in bold.

| Methods | random split |  | leave-one-batch |  | leave-one-cell-type |  |
| --- | --- | --- | --- | --- | --- | --- |
|  | Pearson | Spearman | Pearson | Spearman | Pearson | Spearman |
| moETM | **0.58** (0.03) | **0.33** (0.02) | **0.55** (0.04) | **0.30** (0.03) | **0.51** (0.03) | **0.28** (0.04) |
| BABEL | 0.38 (0.02) | 0.27 (0.03) | 0.34 (0.03) | 0.23 (0.02) | 0.31 (0.03) | 0.18 (0.03) |
| scMM | 0.40 (0.03) | 0.29 (0.03) | 0.37 (0.04) | 0.25 (0.03) | 0.33 (0.03) | 0.21 (0.03) |

**Table S9:** Imputing gene expression from surface protein expression using the BMMC2 dataset. The best score per evaluation metric is in bold.

| Methods | random split |  | leave-one-batch |  | leave-one-cell-type |  |
| --- | --- | --- | --- | --- | --- | --- |
|  | Pearson | Spearman | Pearson | Spearman | Pearson | Spearman |
| moETM | **0.65** (0.03) | **0.41** (0.03) | **0.63** (0.02) | **0.39** (0.03) | **0.60** (0.03) | **0.37** (0.02) |
| BABEL | 0.62 (0.03) | 0.39 (0.03) | 0.60 (0.04) | 0.37 (0.03) | 0.57 (0.02) | 0.33 (0.03) |
| scMM | 0.61 (0.02) | 0.37 (0.03) | 0.60 (0.03) | 0.36 (0.04) | 0.54 (0.04) | 0.30 (0.03) |

### S2 Supplementary Figures

**Figure S1:**
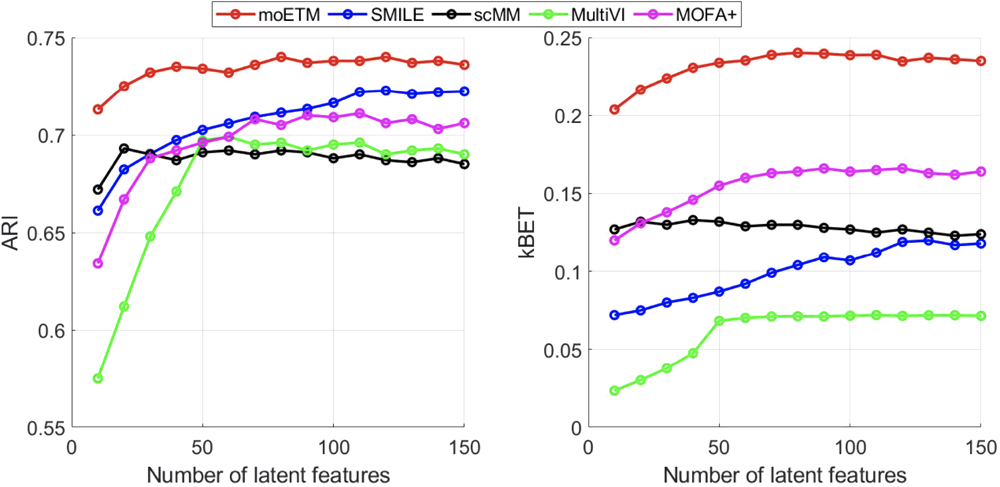
Evaluation metric values as a function of the numbers of latent dimensions on the BMMC1 dataset.

**Figure S2:**
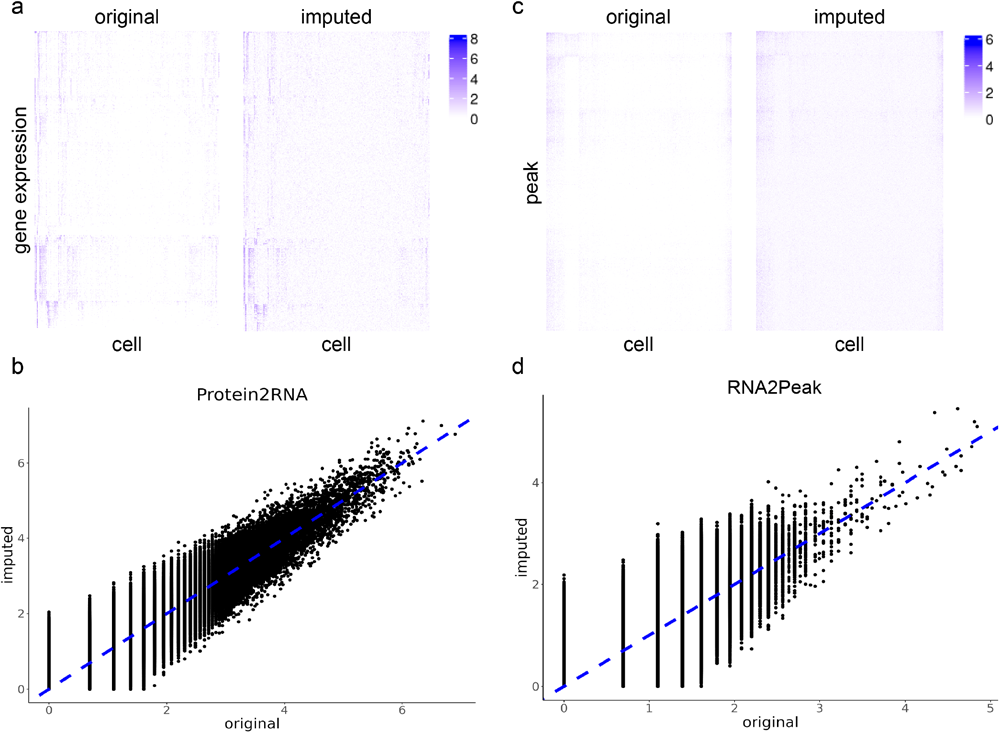
Cross-omic imputation from low dimension to high dimension. **a**. Heatmap of original and imputed gene expression from protein values using the BMMC2 CITE-seq dataset. In each heatmap, the columns are the randomly sampled 5000 cells, and the rows are genes. The order of cells and genes are the same in the two heatmaps. **b**. Scatter plot of original and imputed gene expression values. The x-axis is the original expression and the y-axis represents the imputed values. The diagonal line is in blue. **c**. Heatmap of the original and imputed peak values from gene expression on the BMMC1 dataset. Rows are peaks and columns are randomly sampled 5000 cells. **d**. Scatter plot of the original and imputed peak values.

**Figure S3:**
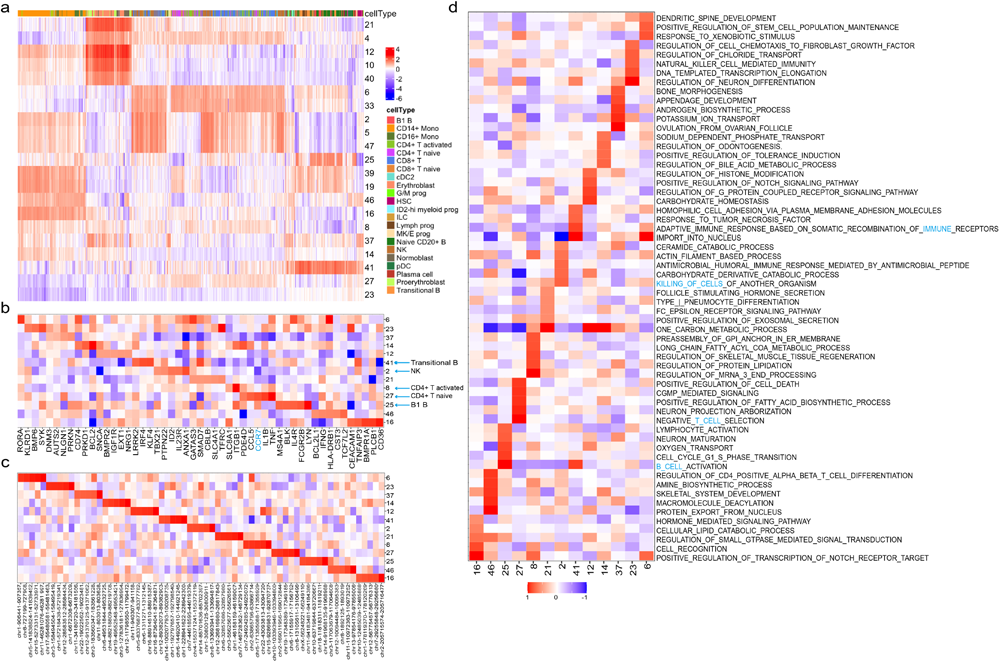
GO-informed cell and features embedding learned by our moETM on the BMMC1 dataset. The gene embedding matrix of moETM was fixed to the gene sets of Gene Ontology Biological Processes from MSigDB during the training on the BMMC1 gene+peak data. **a**. Cell topic-embedding. Columns are cells and rows are topics. The top bar indicate the cell types. Color intensities are proportional to the topic embedding of the cells. **b**. Top genes for the select topics. Rows are the select topics and columns are the top 5 genes per topic. Marker genes were colored in blue. Cell-type-specific topics were labeled by the arrows. **c**. Top peaks for the same set of topics as in panel b. **d**. The top 5 pathways for each of the selected topics. Cell-type-associated pathways were colored in blue. The color intensities are the topic embedding values for the pathways.

**Figure S4:**
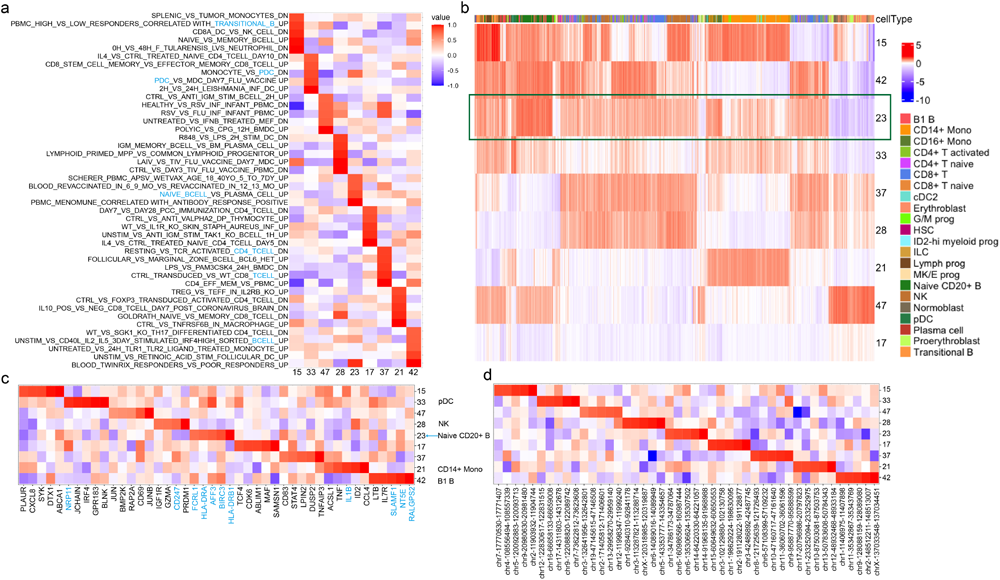
GO-informed cell and features embedding learned by our moETM on the BMMC1 dataset. **a**. The top 5 pathways for each of the selected topics. Cell-type-associated pathways were colored in blue. The color intensities are the topic embedding values for the pathways. **b**. Cell topic-embedding. Columns are cells and rows are topics. The top bar indicate the cell types. Color intensities are proportional to the topic embedding of the cells. The highlighted topic 23 was discussed in the main text. **c**. Top genes for the select topics. Rows are the select topics and columns are the top 5 genes per topic. Marker genes were colored in blue. Cell-type-specific topics were labeled by the arrows. **d**. The top 5 pathways for each of the selected topics. Cell-type-associated pathways were colored in blue. The color intensities are the topic embedding values for the pathways.

**Figure S5:**
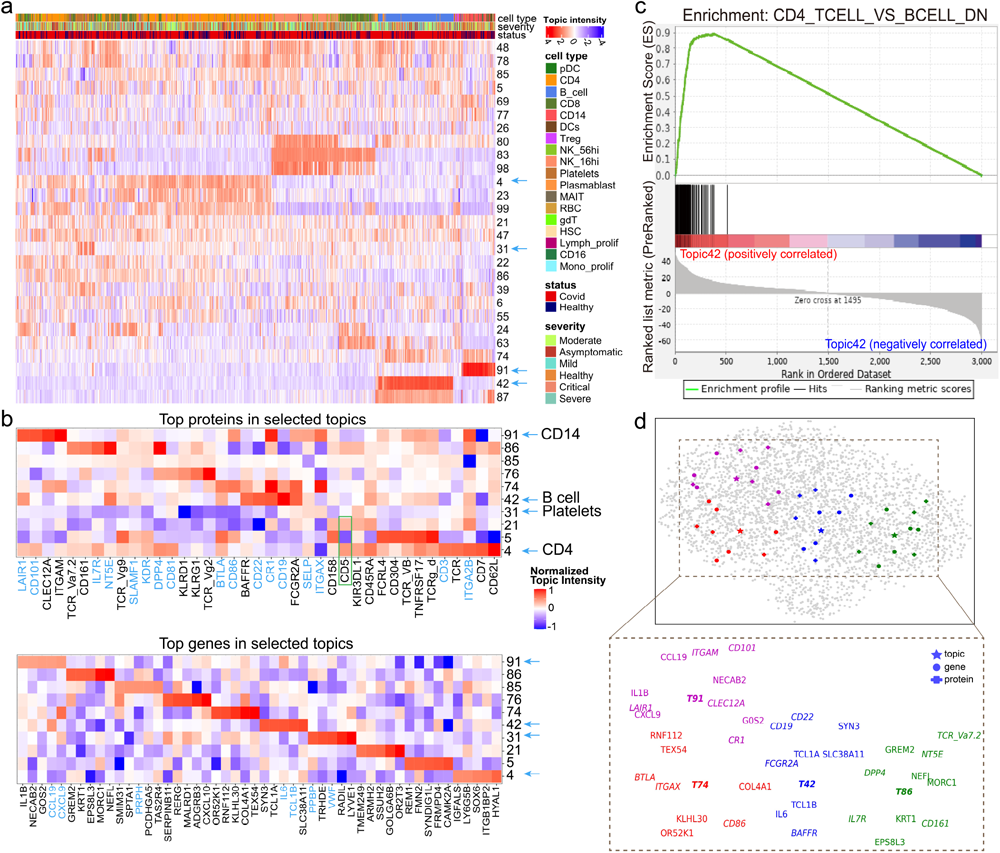
Topic analysis of single-cell COVID-19 CITE-seq dataset. **a**. Topics intensity of cells sub-sampled (n=10000). Only the topics with the sum of absolute values larger than the third quartile across all sampled cells were shown. The three color bars show cell types, disease severity, and disease status. **b**. Top proteins and top genes per select topic. The marker genes and proteins based on CellMarker or literature search are colored in blue. For visualization purposes, we divided the topic values by the maximum absolute value within the same topic. **c**. GSEA lead-edge analysis of topic 42. The enriched gene set contains genes that were down-regulated in CD4 T cells relative to B cells. The barcode in the middle are the genes that belong to the corresponding gene set. **d**. UMAP visualization of the genes, proteins, and topics via their shared embedding space. Genes, proteins, and topics were labeled by star, circle and cross shapes on the top panel, respectively. Topics 42, 74, 86, and 91 were colored in blue, red, green, and purple, respectively. The bottom panel displays the corresponding topic indices and gene symbols highlighted on the top panel.

**Figure S6:**
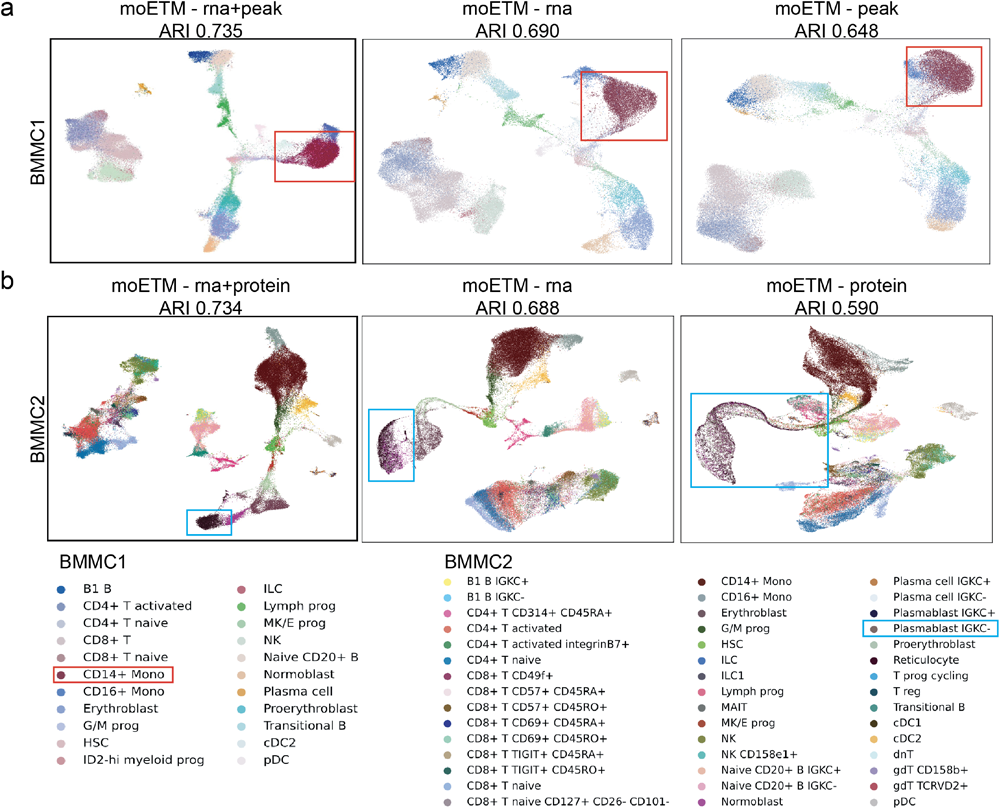
UMAP visualization of single modality embedding and multiple modalities embedding. **a**. UMAP visualizations of individual and integrated embedding in the BMMC1 dataset. Each point represents a cell and different color represents different cell types. The first column is the multiple modalities joint embedding visualization by moETM. The second and third columns are single modality embeddings by moETM_rna/peak. The highlighted clusters and cell types in the legend were described in the main text. **b**. UMAP visualizations of individual and integrated embedding in the BMMC2 dataset.

**Figure S7:**
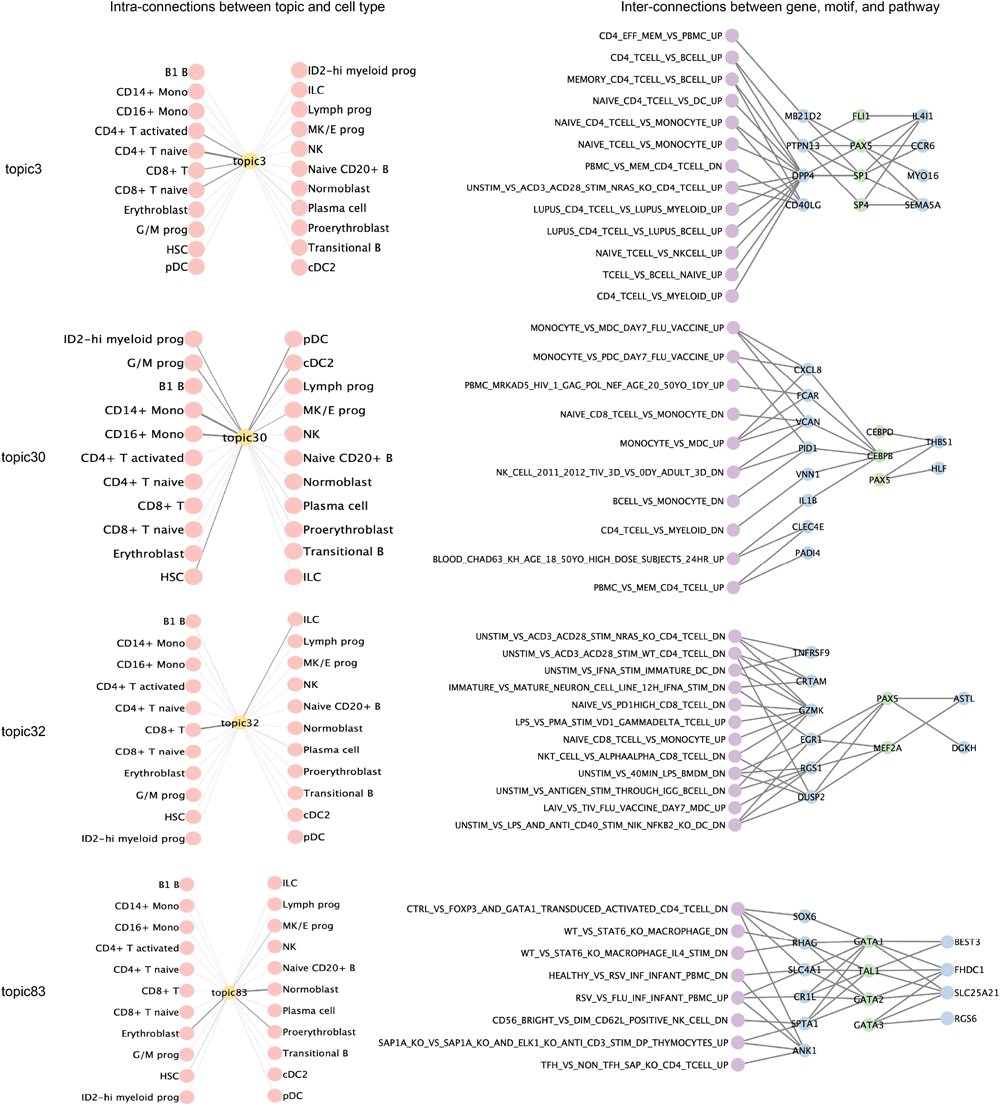
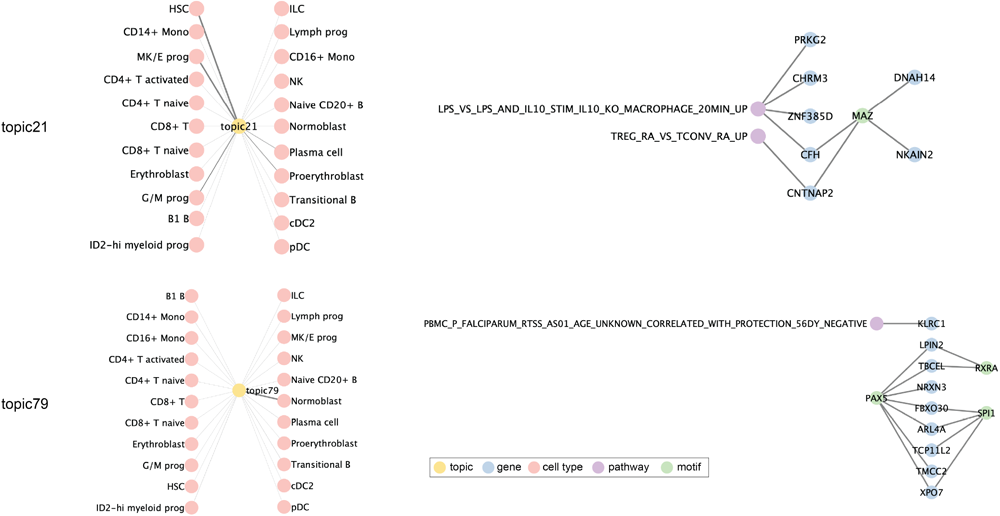
Topic-directed regulatory networks. Topic-directed regulatory networks. Each network contains nodes representing a topic, top genes, enriched pathways, cell types, and enriched motifs under the topic. Different types of nodes are colored in different colors. The left panel represents connections between the topic and cell types, where the edge width is proportional to topic scores. The right panel represents TF-target gene associations and pathway-gene relationships based on ENCODE TF Targets database and the MSigDB database respectively. The edge width for the topic-cell_type is proportional to the enrichment scores. In particular, the most enriched cell types for a topic would contribute to the largest edge width among all topic-cell type edges. For example, topic 32 is the most enriched for CD8+ T cell, and the edge width for topic32-CD8+ T is the largest compared with other cell types. Similarly, the most enriched cell types for topic 79 and topic 21 are normoblast and HSC, respectively, which are indicated by the largest width of the corresponding edges.

**Figure S8:**
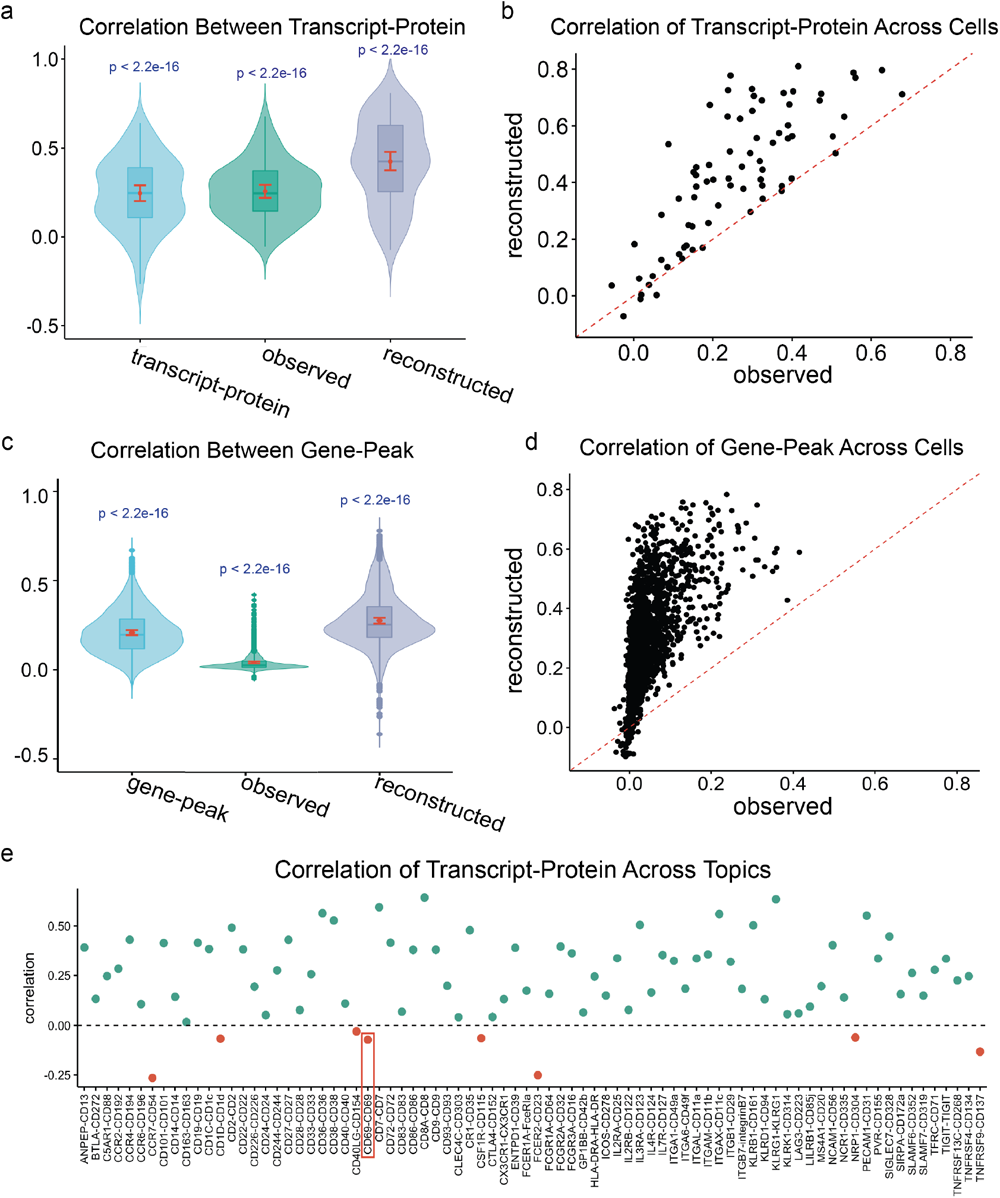
Spearman correlation analysis. **a**. Correlations between transcript-protein across topic scores, observed sample values, and reconstructed sample values in the BMM2 dataset. For each violin plot, a kernel density estimation of the correlations is shown on each side of the gray line. Within the violin plot, the red line indicates the 95% confidence interval of those correlation values as 1.96 times the standard deviation of the correlation scores, and the lower and upper whiskers of boxplot are 25th and 75th quartiles, respectively. The p-values indicated on the top of the violin plots were based on one-sample t-test. **b**. Comparison between geneprotein across sample values in the BMMC2 dataset. The x-axis is the correlation based on observed values and the y-axis is the correlation based on reconstructed values. The red dashed line represents the diagonal. **c,d**. Correlations between gene-peak in the BMMC1 dataset. **e**. Correlations between genes and proteins across topics in the BMMC2 dataset. Each dot represents a gene-protein pair. Negative correlations are colored in red and positive correlations are colored in green. The highlighted pair CD69-CD69 was described in the main text.

**Figure S9:**
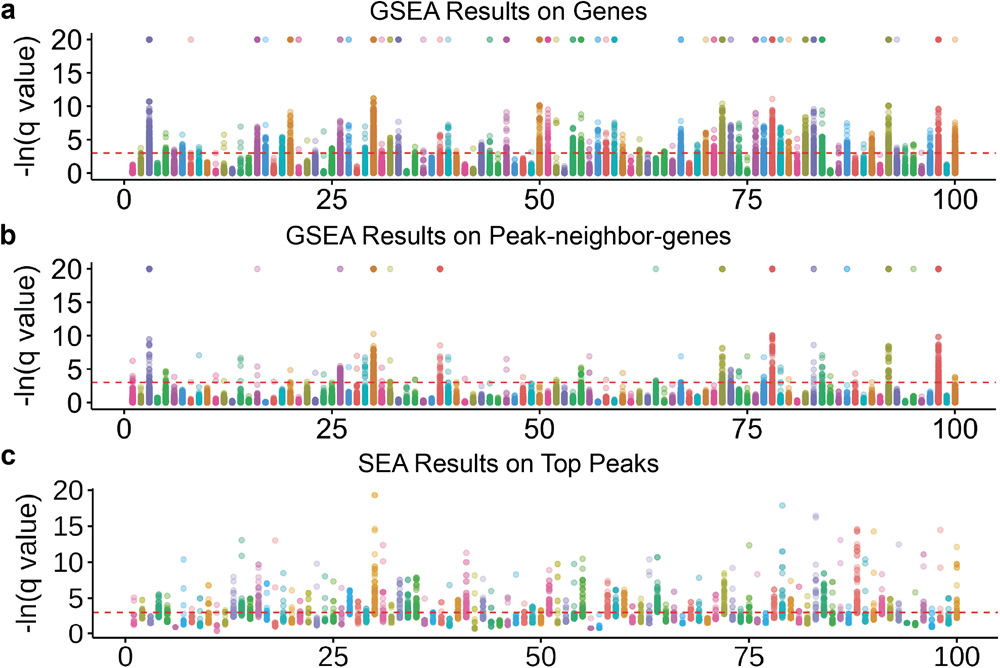
Pathway enrichment and motif enrichment for the 100 topics learned from the BMMC1 data. In each plot, the x-axis is the 100 topics and the y-axis is the pathway enrichment scores or the motif enrichment scores in terms of -ln q-value (i.e., p-values adjusted for multiple testing by Bonferroni correction). The top and middle panels are the corresponding GSEA enrichments of gene and peak-neighbor-gene topic scores, respectively. The bottom panel is the corresponding SEA enrichment of the top 100 peaks.

